# Intrinsic DNA motif-binding capability of OsCLSY3 guides female-preferential RNA-directed DNA methylation

**DOI:** 10.1101/2024.12.03.626448

**Authors:** Taehoon Kim, Marcio F. R. Resende, Meixia Zhao, Kevin Begcy

## Abstract

DNA methylation is essential for regulating gene expression and transposon silencing. In plants, RNA-directed DNA methylation (RdDM) mediates de novo DNA methylation. In *Arabidopsis*, RdDM is targeted to specific loci through either histone methylation or DNA motifs, the latter involving the chromatin remodelers CLASSY3 and CLASSY4 (CLSY3/4) in reproductive tissues mediated through the interaction of REPRODUCTIVE MERISTEM (REM) transcription factors and the GENETICS DETERMINES EPIGENETICS1 (GDE1) linker protein with an AT-rich DNA motif. Here, we identify a transcription factor-independent mechanism for female-preferential RdDM in rice, in which OsCLSY3 directly recognizes a GC-rich DNA motif, termed the Female-preferential RdDM-associated Element (FRE). OsCLSY3’s intrinsic motif-binding capability is distinct from CLSY proteins in *Arabidopsis*. Interaction between OsCLSY3 and FRE-associated loci produces abundant 24-nucleotide small interfering RNAs in female reproductive tissues, driving progressive CHH methylation during development. These findings uncover a rice-specific mode of RdDM and provide insight into conserved and divergent regulation of DNA methylation across plant lineages.

## Main

Cytosine DNA methylation is a key epigenetic modification that maintains genome stability and regulates gene activity^1,2^. Unlike mammals, in which DNA methylation occurs predominantly in the CG context, plants display DNA methylation in three sequence contexts: CG, CHG, and CHH, where H = A, T, or C^1^. Dynamic DNA methylation patterns across developmental transitions are established by the plant-specific RNA-directed DNA methylation (RdDM) pathway^3,4^. Canonical RdDM produces 24-nucleotide small interfering RNAs (24-nt siRNAs) to guide cytosine methylation in all sequence contexts^3,4^, in which the plant-specific RNA polymerase IV (Pol IV) generates single-stranded RNAs, which are converted into double-stranded RNAs by RNA-DEPENDENT RNA POLYMERASE 2 (RDR2) and subsequently cleaved into 24-nt fragments by DICER-LIKE 3 (DCL3)^5–7^. These 24-nt siRNAs are loaded onto ARGONAUTE 4 (AGO4)-family proteins and bind complementarily to Pol V-transcribed scaffold RNAs^8–10^. This leads to the recruitment of DOMAIN REARRANGED METHYLTRANSFERASE 2 (DRM2), which methylates cytosines at nearby loci^11^. Non-canonical RdDM pathways use byproducts of post-transcriptional gene silencing or Pol II-transcribed RNA hairpins, which are processed by RDR6 and/or DCL1/2/4^3,12^.

RdDM-driven CHH methylation is an important epigenetic mark during pre- and post-fertilization reproductive development in *Arabidopsis*^13^, *Brassica rapa*^14^, *Capsella*^15,16^, tomato^17^, maize^18,19^, and rice^20–26^. Unlike *Arabidopsis*, in which most mutations in RdDM genes cause no remarkable developmental defects, in rice, most RdDM mutants exhibit severe fertility defects^20–26^. For example, loss-of-function mutation of the RDR2 ortholog *OsRDR2* results in severe downregulation of 24-nt siRNA expression and CHH methylation during reproductive development, leading to defects in reproductive organs and complete sterility^20^. Tissue-specific regulation of RdDM has been shown to shape the DNA methylation landscape during reproduction. Notably, local enrichment of 24-nt siRNAs was reported in rice endosperm within genomic regions. These abundant enrichments of 24-nt siRNAs were named *siren* loci (<u>siR</u>NA in <u>en</u>dosperm)^27,28^. Similar enrichment was observed in ovules and endosperm in *A. thaliana*, *B. rapa,* and *Capsella*, at the same genomic regions, and were also named *siren* loci^16,29,30^. Unlike these Brassicaceae species, rice shows distinct siRNA patterns between female reproductive organs and endosperm^31,32^, indicating divergence in tissue-specific RdDM across plant lineages and further raising the question of how reproductive RdDM is targeted to distinct genomic regions in different species.

Major determinants of tissue-specific RdDM targeting are the CLASSY (CLSY) chromatin remodeler proteins, which regulate RdDM patterns by recruiting Pol IV to specific genomic regions^33,34^. In *Arabidopsis*, CLSY1/2 target RdDM sites through interaction with SAWADEE DOMAIN HOMOLOG 1 (SHH1), which recognizes dimethylated H3K9 and unmethylated H3K4^33,35–37^. In contrast, CLSY3/4 recruit Pol IV in a histone methylation-independent manner through an AT-rich DNA motif. Recent studies revealed that several REPRODUCTIVE MERISTEM (REM) transcription factors bind to this motif, and that GENETICS DETERMINES EPIGENETICS 1 (GDE1) protein bridges between REM proteins and CLSY3/4^38,39^. In particular, CLSY3 shapes RdDM dynamics at ovule *siren* loci, making it of particular interest with regard to female-specific siRNA accumulation across plant lineages^29,38,39^. Its involvement in *B. rapa* ovule *siren* loci has been inferred from an analogous female-enriched expression pattern and the presence of AT-rich motifs within these loci^40^.

Therefore, known CLSY-mediated targeting mechanisms rely on partner proteins that recognize chromatin marks or DNA motifs, and whether CLSY proteins themselves can directly recognize DNA sequence has remained unexplored.

Here, we identify a specific mechanism for female-preferential RdDM associated with the intrinsic motif-binding activity of OsCLSY3. We define female gametophyte *siren* (FG-*siren*) loci as high-output 24-nt siRNA-producing regions that dominate the small RNA landscape during female reproductive development and undergo progressive CHH methylation. We further identify a Female-preferential RdDM-associated Element (FRE), a GC-rich motif enriched at FG-*siren* loci and show that OsCLSY3 directly interacts with FRE-containing DNA. These findings reveal an autonomous motif-binding capability of a plant CLSY protein and uncover a transcription-factor-independent mechanism for reproductive RdDM targeting in rice.

## Results

### Developmental regulation of RdDM at FG-siren loci

To identify temporal epigenomic dynamics during rice female reproduction, we captured small RNAs (Supplementary Fig. 1 and Supplementary Data 1) and DNA methylation (Supplementary Data 2) profiles from ovaries across six developmental stages spanning the entire trajectory of female gametophyte (FG) development: megaspore mother cell (MMC), meiosis (MEI), early and late mitosis (MIT1–2), and early and late mature embryo sac (MAT1– 2), as previously defined^41^. Flag leaves (FL) were analyzed in parallel as a vegetative tissue control at the same plant age.

To assess whether the sRNA profiles in ovaries during FG development are preset at the beginning of FG development, 24-nt siRNA clusters were identified from individual developmental samples. Among these clusters, we defined 24-nt siRNA abundant loci based on a cumulative sum analysis. Consistent with previous reports of 24-nt siRNA enrichment in a limited number of genomic regions on female reproductive tissues in rice and Brassicaceae species^29–32^, we found that 166 loci accounted for 90.9% of all 24-nt siRNAs expressed in the mature ovary (MAT2 stage) (Fig. 1a and Supplementary Data 4). Of these, nearly all (95.8%) were common 24-nt siRNA abundant loci across all developmental stages (Fig. 1b and Supplementary Fig. 2a–c). Some of these common loci were physically located near each other (Fig. 1c and Supplementary Fig. 2d). Thus, after merging adjacent loci within 1 kb, we obtained a final set of 137 common 24-nt siRNA loci (Fig. 1d and Supplementary Fig. 2e). In rice endosperm, a small number of genomic regions dominate 24-nt siRNA production and are termed “*siren*” (<u>siR</u>NA in <u>en</u>dosperm) loci^27,28^. Similar loci have been described in Brassicaceae ovules, where the same genomic regions produce 24-nt siRNAs in both ovules and endosperm^29,30^. In contrast, 24-nt siRNA-expressing loci in rice female reproductive tissues showed limited overlap with those in rice endosperm^32,42^ (Supplementary Fig. 7). Therefore, we termed these 24-nt siRNA abundant loci as “FG-*siren*”, reflecting their similarity to dicot ovules^29,30^, while distinguishing them from rice endosperm *siren* loci^27,28^.

**Fig. 1.**
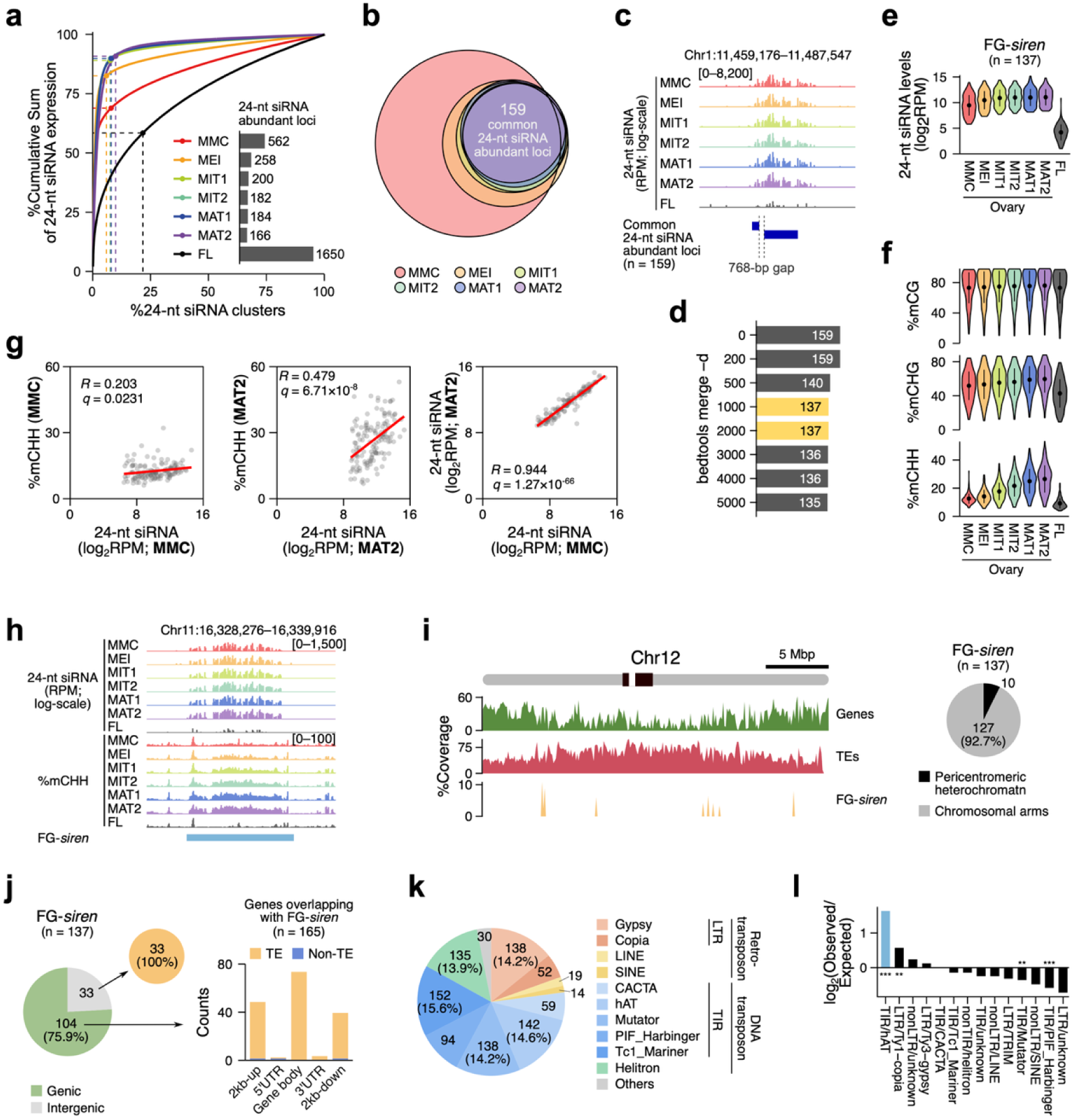
Identification and characterization of FG-*siren* loci. **a,** Relationship between the percentage of 24-nt siRNA clusters (*x* axis) and the cumulative sum of RPM-normalized expression (*y* axis). 24-nt siRNA abundant loci were identified at the knee point of each graph. Inset bar charts indicate the number of 24-nt siRNA abundant loci identified from each sample. Genomic coordinates of identified loci are listed in Supplementary Data 4. **b,** Overlap among 24-nt siRNA abundant loci identified from ovaries at each stage (MMC–MAT2). **c,** Representative FG-*siren* locus generated by merging two adjacent loci. **d,** Number of FG-*siren* loci obtained at each merge-gap threshold; the selected threshold and resulting final set of FG-*siren* loci (137 loci) are highlighted in yellow. Genomic coordinates of FG-*siren* loci are listed in Supplementary Data 5. **e,f,** 24-nt siRNA expression (**e**) and DNA methylation levels (**f**) at FG-*siren* loci. Dots and lines denote medians and interquartile ranges, respectively. **g,** Correlation between 24-nt siRNA and CHH methylation levels in ovaries at MMC and MAT2 stages. Red lines represent simple linear regression fits. Pearson correlation coefficient (*R*) and Benjamini-Hochberg (BH)-adjusted false discovery rates (*q*) are indicated. **h,** 24-nt siRNA and CHH methylation levels at a representative FG-*siren* locus. **i,** Chromosomal distribution of genes, TEs, and FG-*siren* loci along chromosome 12, summarized in 100-kb bins (left panel). Pie chart showing genome-wide distribution of FG-*siren* loci (right panel). Pericentromeric heterochromatin was defined based on the genetic recombination-suppressed regions reported by Tian et al.^75^; the remaining regions were considered chromosomal arms. **j,** Count of genomic features overlapping with FG-*siren* loci. The number of TE and non-TE elements with respect to the genic elements are shown. **k,** Proportion of TE families overlapping with FG-*siren* loci. **l,** Observed overlap of TE families with FG-*siren* loci compared to the expected overlap, calculated based on their genomic proportions. Statistical significance was determined using a two-tailed Fisher’s exact test (\*\*\**P* < 0.001; \*\**P* < 0.01). Overrepresented TE families (observed/expected > 2; *P* < 0.05) are highlighted in light blue.

To investigate whether 24-nt siRNAs generated at FG-*siren* loci direct DNA methylation during FG development, as established for RdDM^3^, we examined the distribution of 24-nt siRNA expression and DNA methylation across these regions (Fig. 1e–h and Supplementary Figs. 2–4). We found that all FG-*siren* loci exhibited high levels of 24-nt siRNAs in ovaries, with an increasing trend during the FG development, whereas flag leaves showed only marginal expression (Fig. 1e,h). The 24-nt siRNA landscape in flag leaves was distinct from that in ovaries (Supplementary Fig. 3). CHH methylation in these regions increased progressively, from a median of 12.0% at MMC to 27.0% at MAT2, whereas CHG methylation increased more modestly (from 51.5% to 59.4%) in the same developmental window. In contrast, flag leaves showed reduced levels of CHG and CHH methylation relative to developing ovaries. CG methylation remained largely unchanged both across FG developmental stages in ovaries and between ovaries and flag leaves (Fig. 1f,h). These results suggest that RdDM at FG-*siren* loci is preferential in female reproductive tissues and developmentally reinforced, consistently with previous reports indicating that 24-nt siRNAs primarily influence CHH methylation during rice reproduction, leading to overall elevated CHH methylation levels in reproductive tissues compared to vegetative tissues^20^. Interestingly, 24-nt siRNA levels were moderately correlated with CHH methylation at the MMC stage (*R* = 0.203) but showed a stronger positive correlation at the MAT2 stage (*R* = 0.479), despite highly similar siRNA levels between these stages (*R* = 0.944) (Fig. 1g,h and Supplementary Fig. 4). This suggests that 24-nt siRNA accumulation at FG-*siren* loci is established in earlier stages of the female reproductive lineage, whereas CHH methylation is progressively established during FG development, consistent with developmental time-lag in RdDM during *Arabidopsis* embryogenesis^43^.

Next, we examined the chromosomal distribution of FG-*siren* loci. Most FG-*siren* loci (92.7%) were enriched in the chromosomal arms, while only 7.3% were located in pericentromeric heterochromatin (Fig. 1i and Supplementary Fig. 14). In contrast to Brassicaceae, where ovule *siren* loci are evenly distributed across chromosomal regions^29,30^, this distribution suggests evolutionary divergence in the mechanisms underlying locus-specific 24-nt siRNA accumulation between monocots and dicots. More than 75% of the FG-*siren* loci overlapped with genic regions. Of these, nearly all (98.2%) overlapped with TEs on gene bodies or 2 kb flanking regions of genes. All FG-*siren* loci located in intergenic regions also overlapped with TEs (Fig. 1j). Among TEs overlapping with FG-*siren* loci, DNA transposons accounted for 74.0%, of which hAT, Mutator, and Tc1-Mariner being the most abundant subfamilies (Fig. 1k). However, only hAT elements were significantly overrepresented relative to their genome-wide abundance (Fig. 1l). Together, these results suggest that FG-*siren* loci represent TE-associated siRNA-producing regions with developmentally regulated activity during rice FG development.

### Female-preferential RdDM at FG-siren loci

Given the high 24-nt siRNA levels and CHH methylation at FG-*siren* loci in developing ovaries but not in flag leaves (Fig. 1e,f,h and Supplementary Fig. 3), we compared siRNA and DNA methylation profiles across vegetative and reproductive tissues from different rice cultivars to assess the tissue specificity of RdDM at these regions^20–24,26–28,31,32,42,44–55^ (Supplementary Data 1,2). FG-*siren*-derived 24-nt siRNAs were nearly undetectable in vegetative tissues and remained low in male reproductive tissues despite their overlap with 24-nt siRNA clusters in seedlings (Fig. 2a,c), resulting in approximately 10% of CHH methylation levels in these organs (Fig. 2b). In contrast, female reproductive tissues showed strong enrichment of 24-nt siRNAs throughout development, accompanied by CHH methylation, with ∼70% of 24-nt siRNA reads mapping to FG-*siren* loci (∼0.23% genome coverage) in mature ovaries and pistils (Fig. 2a,b,e). This imbalance between male and female reproductive tissues suggests that the overall siRNA abundance in mature panicles is largely driven by female reproductive tissues (Fig. 2a,c–e and Supplementary Fig. 5a). Notably, this enrichment was also observed in gametic eggs and zygotes (9 hours after pollination), despite its absence in the sperm cells (Fig. 2a and Supplementary Fig. 5a), indicating that FG-*siren* loci largely shape the sRNA landscape across pre- and post-fertilized stages along the female lineage. However, the embryo did not show such enrichment (Fig. 2a and Supplementary Fig. 5a), indicating that 24-nt siRNA targeting is likely reprogrammed during embryogenesis, consistent with dynamic changes in the sRNA landscape during *Arabidopsis* embryo development^43,56^.

**Fig. 2.**
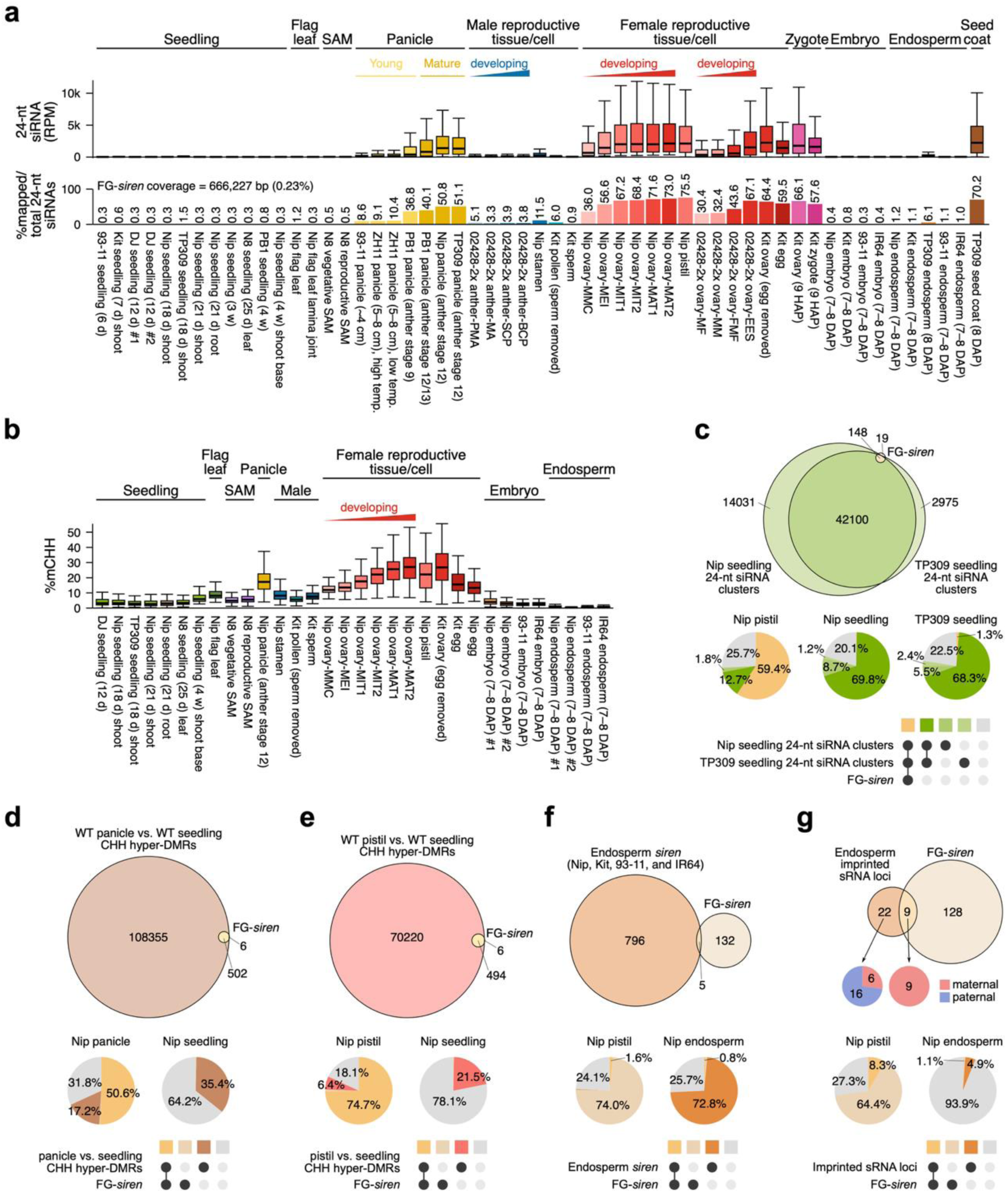
Female-preferential RdDM at FG-*siren* loci. **a,b,** 24-nt siRNA expression (**a**) and CHH methylation levels (**b**) at FG-*siren* loci across vegetative and reproductive tissues from different rice varieties. In the lower panel of **a**, the relative abundance of 24-nt siRNAs derived from FG-*siren* loci is shown. sRNA-seq and BS-seq datasets obtained from public repositories^20–24,26–28,31,32,42,44–55^ and are listed in Supplementary Data 1 and 2, respectively. **c,** Comparison of FG-*siren* loci with seedling 24-nt siRNA clusters. **d,e,** Comparison between FG-*siren* loci and CHH hyper-DMRs in Nipponbare panicles (**d**) and pistils (**e**) compared to seedlings. **f,** Comparison between FG-*siren* loci and endosperm *siren* loci. **g,** Comparison between FG-*siren* loci and endosperm imprinted sRNA loci. For **c–g,** scaled Venn diagrams depict the relationships among the compared loci, and pie charts show the relative abundance of 24-nt siRNAs in each intersection category. For **g,** inset pie charts show the proportion of maternally and paternally imprinted sRNA loci in each intersection category. Genome coordinates for seedling 24-nt siRNA clusters^23^, CHH hyper-DMRs in panicles and pistils compared to seedlings^20^, endosperm *siren* loci^28^, and endosperm imprinted sRNA loci^27^ were obtained from previous studies, and comparisons with FG-*siren* loci are listed in Supplementary Data 7–9. Abbreviations for rice cultivars: Nip, Nipponbare; Kit, Kitaake; TP309, Taipei 309; ZH11, Zhonghua 11; N8, Norin 8; DJ, Dongjin; PB1, Pusa Basmati 1.

### Distinct RdDM regulation between female reproductive tissues and endosperm in rice

Although concentrated 24-nt siRNA expression in rice was first described at endosperm *siren* loci^27,28^, 24-nt siRNA expression at FG-*siren* loci is largely absent in endosperm (Fig. 2a,f and Supplementary Figs. 6h,7). Endosperm *siren* loci showed only limited overlap with FG-*siren* loci, although ≥70% of 24-nt siRNAs in endosperm and pistils mapped to endosperm *siren* and FG-*siren* loci, respectively (Fig. 2f and Supplementary Fig. 7). Seed coat, derived from somatic integuments of ovaries, showed higher 24-nt siRNA levels at FG-*siren* loci, comparable to mature ovaries and pistils (Fig. 2a and Supplementary Fig. 6i). In Brassicaceae including *Arabidopsis*, *Brassica*, and *Capsella*, endosperm siRNAs are thought to arise mainly through non-cell-autonomous movement of maternally expressed siRNAs from the seed coats^16,30^.

However, a similar phenomenon has not been demonstrated in rice. Interestingly, 9 out of 15 maternally imprinted siRNA loci overlapped with FG-*siren* loci, whereas none of the paternal counterparts did. Nonetheless, the proportion of maternally derived siRNAs in rice endosperm was marginal (Fig. 2g and Supplementary Fig. 8). Given that rice endosperm *siren* loci are largely distinct from FG-*siren* loci, which show preferential siRNA expression in ovaries, pistils, egg cell, and seed coat (Fig. 2a,f and Supplementary Figs. 6,7), this suggests that the molecular mechanisms underlying tissue-specific RdDM may differ between female reproductive organs and endosperm in rice, as well as how endosperm siRNA landscape is shaped may differ between monocots and dicots.

### FG-siren loci are high-output RDR2-dependent RdDM targets

To determine whether RdDM activity at FG-*siren* loci operates through the canonical pathway, we analyzed 24-nt siRNA accumulation and DNA methylation patterns in *osrdr2* mutants, including the null mutant *osrdr2-3* and the partial-loss-of-function mutant *osrdr2-6*, which exhibit defects in 24-nt siRNA production and CHH methylation in reproductive tissues^20^. At FG-*siren* loci, the high 24-nt siRNA accumulation and CHH methylation observed in reproductive tissues were markedly reduced in *osrdr2* mutants and *osnrpd1* knockdown mutants (Fig. 3a,c–f and Supplementary Fig. 9). Notably, although *osrdr2-6* retained reduced 24-nt siRNA accumulation in these regions, CHH methylation was nearly abolished, similar to *osrdr2-3* (Fig. 3a,d,f), indicating that residual 24-nt siRNA production is insufficient to sustain CHH methylation at these loci. This may reflect a requirement for earlier developmental RdDM activity and/or broader disruption of RdDM capacity in *osrdr2* mutants^20^. Consistent with this, FG-*siren*-derived 24-nt siRNA expression was not impaired in panicles of *osrdr6-1* and *osdcl4-1* knockdown mutants (Fig. 3b), despite 21–22-nt siRNAs exhibiting a spatial distribution similar to that of 24-nt siRNAs at these loci, albeit to a lesser extent (Fig. 2a and Supplementary Fig. 5a). Given that these reproductive 21–22-nt siRNAs are reduced in *osrdr2* mutants and Pol IV-related mutants, but not in *osrdr6-1* and *osdcl4-1* (Supplementary Figs. 9a,10a,d), these results indicate that RdDM at FG-*siren* loci is largely independent of the non-canonical pathway. This further supports that both 24-nt and 21–22-nt siRNAs in these regions are primarily generated through the Pol IV-RDR2 pathway, consistent with previous studies in *Arabidopsis* and *Capsella*^15,57^.

**Fig. 3.**
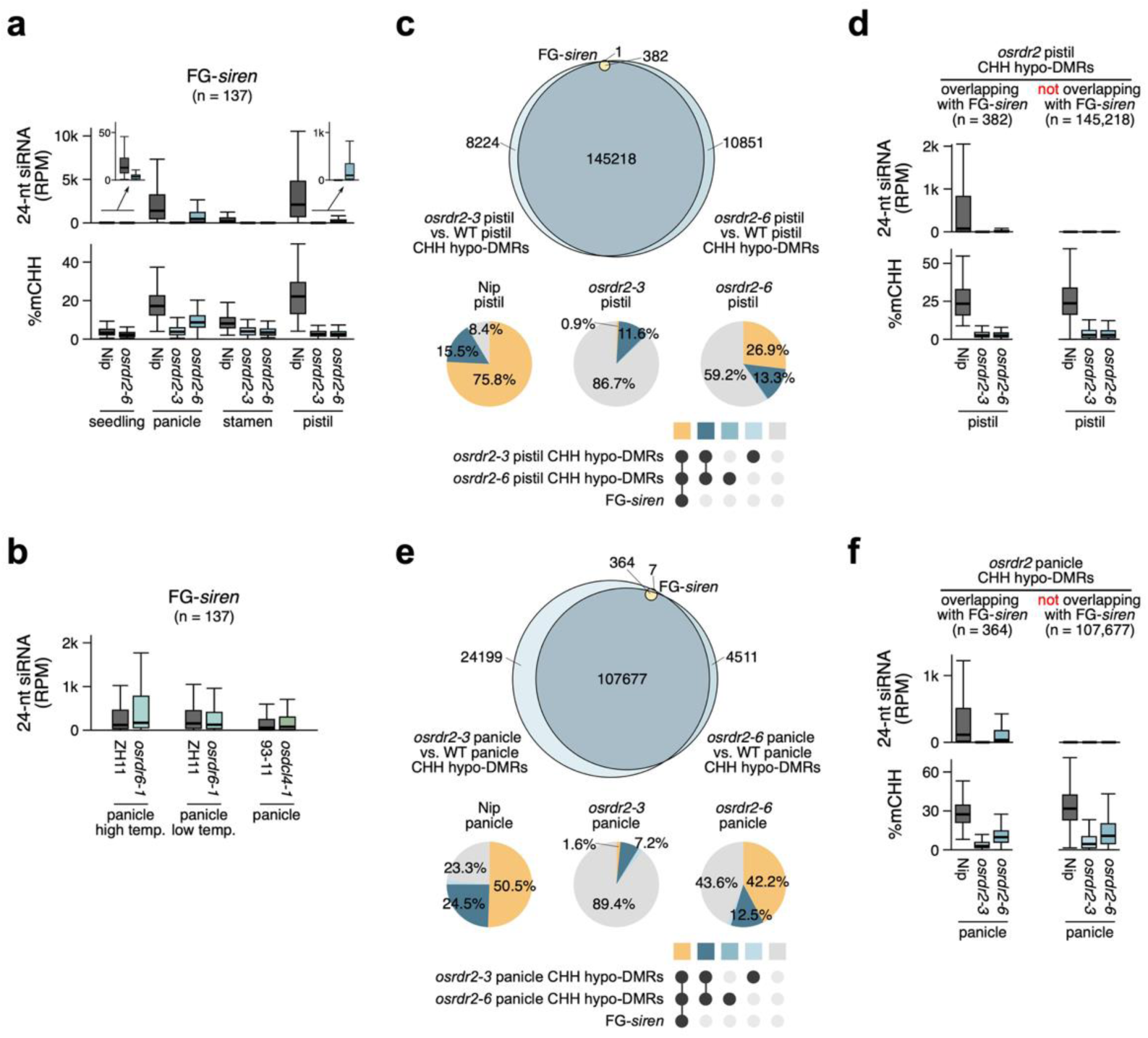
FG-*siren* loci dominate 24-nt siRNA production within RDR2-dependent targets in reproductive tissues. **a,** 24-nt siRNA expression and CHH methylation levels at FG-*siren* loci in WT and *osrdr2* mutants across vegetative and reproductive tissues. **b,** 24-nt siRNA expression levels at FG-*siren* loci in WT, *osrdr6-1*, and *osdcl4-1* mutants in panicles. **c–f,** Comparison between FG-*siren* loci and *osrdr2* CHH hypo-DMRs in pistils (**c,d**) and panicles (**e,f**) compared to WT. For **c,e**, scaled Venn diagrams depict the relationships among the compared loci, and pie charts show the relative abundance of 24-nt siRNAs expressed in each intersection category. For **d,f**, boxplots show 24-nt siRNA expression and CHH methylation levels at *osrdr2* CHH hypo-DMRs categorized by their overlap with FG-*siren* loci. sRNA-seq and BS-seq datasets obtained from public repositories^20,46,47^ and are listed in Supplementary Data 1 and 2, respectively. Genome coordinates for *osrdr2* CHH hypo-DMRs in pistils and panicles compared to WT were obtained from a previous study^20^, and comparisons with FG-*siren* loci are listed in Supplementary Data 10. Abbreviations for rice cultivars: Nip, Nipponbare; ZH11, Zhonghua 11.

Genome-wide loss of RDR2 resulted in widespread CHH hypomethylation in reproductive tissues. However, only a small fraction of hypo-DMRs overlapped with FG-*siren* loci, despite nearly all FG-siren loci being included within these regions (Fig. 3c,e). Consistently, 24-nt siRNA production was highly concentrated at FG-*siren*-overlapping regions, whereas most hypo-DMRs showed little or no siRNA accumulation (Fig. 3c–f). Together, these results suggest that RdDM activity is functionally partitioned, with FG-*siren* loci acting as major sources of 24-nt siRNAs, whereas the majority of RdDM targets represent regions of developmentally regulated methylation with limited siRNA production. Given that OsRDR2-mediated CHH methylation is required for sexual reproduction including normal FG development^20^, these findings further suggest that FG-*siren*-derived siRNAs likely play a critical role in rice reproductive development.

### FRE guides female-preferential RdDM at FG-siren loci

Given that DNA motifs guide Pol IV-mediated RdDM at ovule *siren* loci in *A. thaliana*^29,38,39^, we hypothesized that FG-*siren* loci are regulated by sequence-specific *cis*-elements. We therefore searched for *cis*-regulatory motifs enriched in FG-*siren* loci (Fig. 4, Supplementary Fig. 12, and Supplementary Data 11). Among four identified motifs, Motif1 (GGGGGCMWWKGCCCCC) was palindromic, GC-rich, and highly conserved, occurring in nearly all FG-*siren* loci (97.8%) (Fig. 4a,b). We named this motif FRE (<u>F</u>emale-preferential <u>R</u>dDM-associated <u>E</u>lement). Given that rice genome contains GC-rich chromosomal arms^58^, GC-rich composition of FRE motif may explain why FG-*siren* loci were found mostly in chromosomal arms (Fig. 1i,j and Supplementary Fig. 14). Notably, the FRE motif exhibited a strong positional bias toward the center of FG-*siren* loci (Fig. 4c), and motifs with high similarity to the consensus sequence were preferentially enriched at FG-*siren* loci (Fig. 4d).

**Fig. 4.**
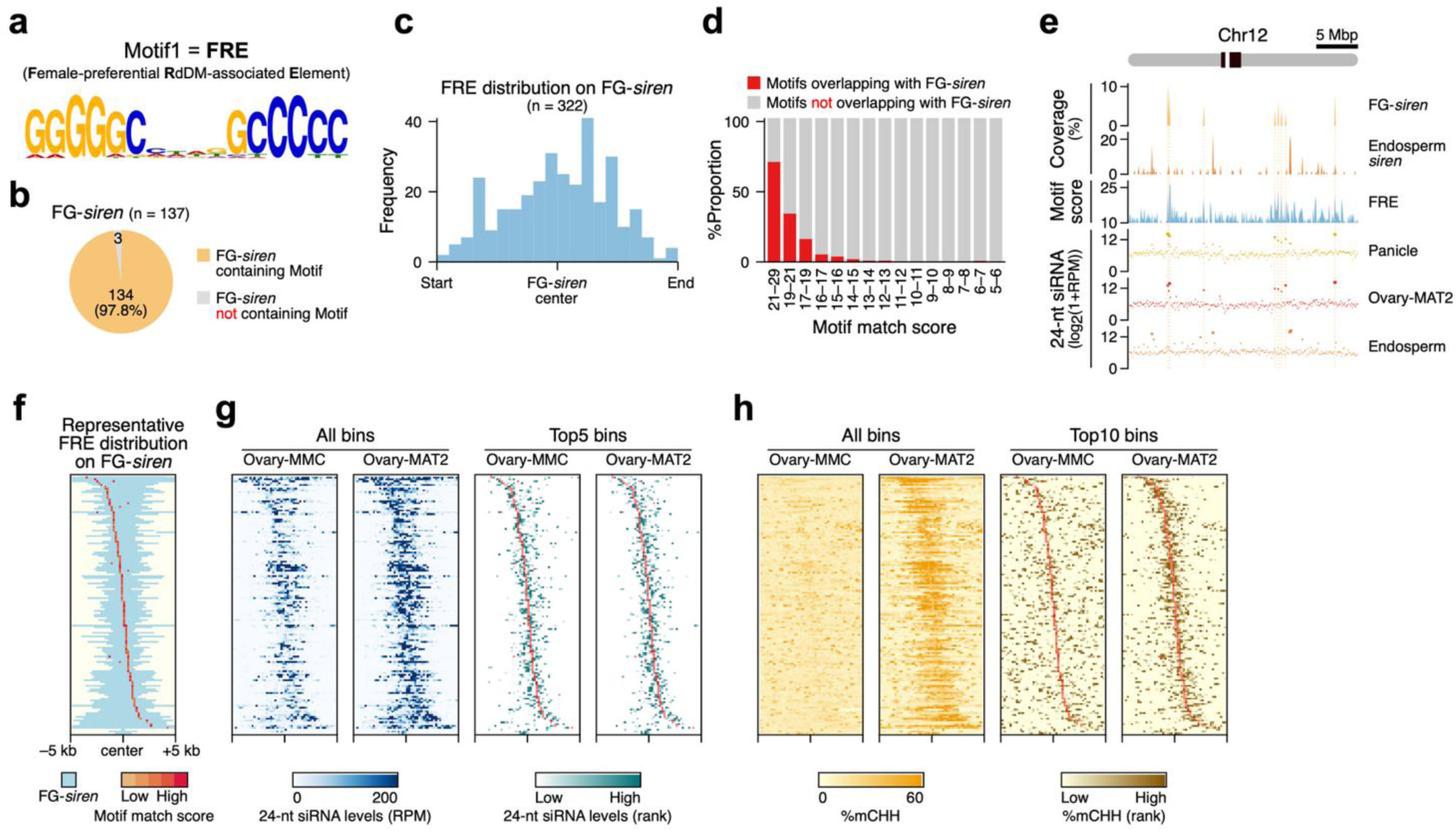
FRE guides female-preferential RdDM at FG-*siren* loci. **a,** DNA motif sequence enriched in FG-*siren* loci. **b,** FRE occurrence at FG-*siren* loci. **c,** Distribution of FRE across FG-*siren* loci in proportional bins (n = 20). **d,** Proportion of motifs overlapping with FG-*siren* loci across motif match score bins, where scores are defined by similarity to the consensus sequence. **e,** Chromosomal distribution of FG-*siren* loci, endosperm *siren* loci, FRE motifs, and 24-nt siRNA levels in Nipponbare panicle, ovary (MAT2 stage), and endosperm along chromosome 12, summarized in 100-kb bins. Pericentromeric heterochromatin was defined based on the genetic recombination-suppressed regions reported by Tian et al.^75^; the remaining regions were considered chromosomal arms. sRNA-seq datasets for Nipponbare panicle and endosperm were obtained from public repositories^20,27^ and are listed in Supplementary Data 1. Genome coordinates for endosperm *siren* loci were obtained from a previous study^28^. **f–h,** Heatmaps showing FRE distribution (**f**), 24-nt siRNA levels (**g**), and CHH methylation levels (**h**) across FG-*siren* loci. For each locus, motifs with the highest match scores were selected as representative. For **g,h**, the top five and top ten bins per locus were highlighted for 24-nt siRNA and CHH methylation levels, respectively.

Further analysis of FRE sequence variants showed that the predominant FRE motifs within FG-*siren* loci motifs retained the terminal GC-rich core sequences and were enriched for A/T bases within the internal region (Supplementary Fig. 13). Furthermore, the FRE motif is located in loci with abundant 24-nt siRNA expression in panicle and ovaries, but not in endosperm (Fig. 4e and Supplementary Fig. 14). Both 24-nt siRNA abundance and CHH methylation peaked at motif-centered regions within individual loci, indicating co-localization of siRNA production and DNA methylation at these sites (Fig. 4f–h). Together, these results indicate that FRE marks sites of female-preferential 24-nt siRNA production and RdDM activity at FG-*siren* loci, suggesting a potential role in directing locus-specific RdDM.

### OsCLSY3: a candidate guiding female-preferential RdDM at FG-siren loci

To identify trans-acting factors that may recognize FRE and mediate RdDM at FG-*siren* loci, we ranked rice proteins based on the correlation between their transcript levels and 24-nt siRNA abundance at FG-*siren* loci across tissues. We integrated previously generated RNA-seq data^41^ with publicly available datasets (Supplementary Data 3), together with corresponding sRNA-seq profiles (Supplementary Data 1) for cross-tissue comparison. Among these, the chromatin remodeler *OsCLSY3* exhibited a strong positive correlation (*R* = 0.514) between its transcript levels and 24-nt siRNA abundance at FG-*siren* loci, representing the highest correlation among DNA methylation-related genes. In contrast, its paralogs *OsCLSY1* and *OsCLSY4* showed negative correlations (Fig. 5a and Supplementary Data 12). Notably, *OsCLSY3* transcript levels were abundant in female reproductive organs, followed by panicles and endosperm^21^, whereas *OsCLSY1* and *OsCLSY4* showed broader expression (Supplementary Fig. 18a,b). Given that *CLSY* expression patterns shape tissue-specific methylation patterns via RdDM^29^ and that knockdown of *OsCLSY3* suppresses RdDM in endosperm^21^, these results suggest a potential role for *OsCLSY3* in female-preferential RdDM at FG-*siren* loci. Given its role in endosperm RdDM^21^, we compared 24-nt siRNA levels in the *osclsy3-kd* mutant. FG-*siren*-derived 24-nt siRNAs were modestly increased in *osclsy3-kd* endosperm compared to wild type (Supplementary Fig. 11a,d), although overall siRNA levels at these loci remain low in endosperm relative to female reproductive tissues (Fig. 2a,f and Supplementary Figs. 7,11).

**Fig. 5.**
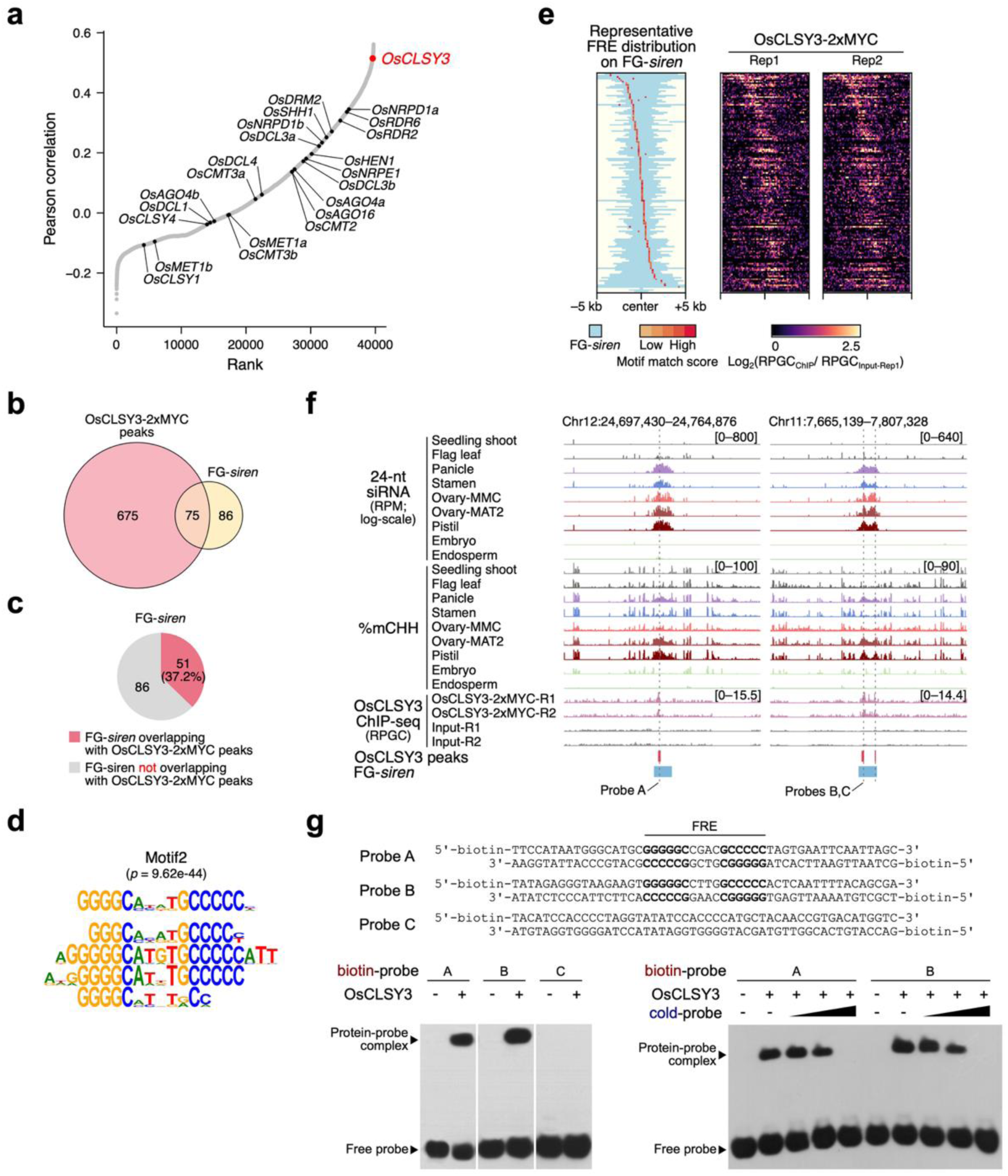
OsCLSY3 directly binds FRE at FG-*siren* loci. **a,** Pearson correlation between 24-nt siRNA levels at FG-*siren* loci and transcript levels of each expressed gene across different rice tissues. Each dot represents a gene. DNA methylation-related genes are highlighted. Detailed information is available in Supplementary Data 11. **b,** Comparison between OsCLSY3-2xMYC ChIP peaks and FG-*siren* loci. **c,** Proportion of FG-*siren* loci overlapping with OsCLSY3-2xMYC ChIP peaks. Identified ChIP peaks and the comparisons with FG-*siren* loci are listed in Supplementary Data 14 and 16, respectively. **d,** FRE-like motifs enriched within OsCLSY3-2xMYC ChIP peaks. Identified motifs are listed in Supplementary Data 15. **e,** Heatmaps showing OsCLSY3 ChIP enrichment across FG-*siren* loci. For each locus, motifs with the highest match score were selected as representative. **f,** 24-nt siRNA expression and CHH methylation levels across rice tissues, together with OsCLSY3 enrichment in rice panicles, at two representative FG-*siren* loci. Grey dotted lines indicate FRE motifs. Mapping statistics of OsCLSY3 ChIP-seq datasets^21^ are listed in Supplementary Data 13. **g,** EMSA showing interactions between truncated OsCLSY3 and FRE-containing probes. Cold-probe gradients represent 7.5-, 15-, and 30-fold excess over biotin-labeled probes. Probe sequences were determined based on the genomic sequences of the representative FG-*siren* loci in **f**. Probes used for EMSA are listed in Supplementary Data 17.

These results likely reflect the reprogramming of RdDM regulation following fertilization^27,32,54^.

### OsCLSY3 directly interacts with FRE for female-preferential RdDM

To determine whether OsCLSY3 is enriched at FRE-containing loci in vivo, we analyzed OsCLSY3 ChIP-seq datasets generated from panicles before anthesis^21^ (Supplementary Note 1, Supplementary Fig. 15, and Supplementary Data 13). Although datasets from two different epitopes (GFP and 2xMYC) were available and their replicates showed high reproducibility (Supplementary Fig. 15a,b), only OsCLSY3-2xMYC ChIP peaks overlapped with FG-*siren* loci (Fig. 5b, Supplementary Figs. 15d, and Supplementary Data 16), which may reflect differences in epitope tagging. Notably, 37.2% of FG-*siren* loci overlapped with ChIP peaks (Fig. 5c, Supplementary Fig. 15d, and Supplementary Data 16), with signals enriched at FRE motifs within each locus (Fig. 5e,f and Supplementary Figs. 15e,f, 16). De novo motif discovery from ChIP peaks identified FRE-like motifs as highly enriched (Fig. 5d, Supplementary Fig. 15c, and Supplementary Data 15). Together, these results suggest that OsCLSY3 is preferentially associated with FRE-containing FG-*siren* loci in vivo and support a role in directing 24-nt siRNA production at these regions.

Although *Arabidopsis* CLSY3 controls RdDM in ovules by interacting with an AT-rich DNA motif through the REM transcription factors and GDE1 linker protein^29,38,39^, our results showed that OsCLSY3 associates with a GC-rich motif (Figs. 4a,5d–f, Supplementary Figs. 15c,e,f,16, and Supplementary Data 15), suggesting that the molecular mechanisms by which CLSY proteins interact with RdDM-associated *cis*-regulatory elements may differ between rice and *Arabidopsis*. To assess the potential for direct interaction between OsCLSY3 and the FRE motif, we used AlphaFold3 (AF3)^59^. To ensure the reliability of multi-molecular predictions, we used a truncated OsCLSY3 (870–1,439) containing the conserved N-terminal SNF2 and C-terminal helicase domains, with high prediction confidences (Supplementary Fig. 17a,b). AF3 predicted a high probability of interaction between the truncated OsCLSY3 protein and FRE-containing probes (0.70 ≤ ipTM ≤ 0.81; 0.80 ≤ pTM ≤ 0.85), but not with non-target probes (Supplementary Fig. 17c–e). To further test this interaction biochemically, we performed electrophoretic mobility shift assays (EMSA). Recombinant OsCLSY3 bound specifically to FRE-containing probes in vitro, despite all probes having a similar GC content (52–56%) (Fig. 5g and Supplementary Fig. 18). Its closest paralog OsCLSY4 did not bind FRE-containing probes (Supplementary Fig. 18), indicating that direct binding to FRE is a unique feature of OsCLSY3. Overall, these results indicate that OsCLSY3’s intrinsic motif-binding capability is distinct from that of *Arabidopsis* CLSY proteins, and that OsCLSY3 may have acquired a novel function during grass evolution, consistent with the previously proposed Poaceae-specific duplication within *CLSY3/4* clades^60^, which led to neofunctionalization of *OsCLSY3* in rice.

## Discussion

In Arabidopsis, CLSY proteins recruit Pol IV through interactions with proteins that recognize either histone methylation or AT-rich DNA motifs^29,33^. Specifically, CLSY3/4 direct reproductive RdDM through the interaction with the GDE1 linker protein and REM transcription factors, which recognize AT-rich DNA motifs^38,39^. However, whether CLSY proteins regulate RdDM with the same molecular mechanisms in other plant species remained unclear. Here, we show that rice CLSY3 guides female-preferential reproductive RdDM at FG-*siren* loci and directly interacts with GC-rich FRE motifs, revealing a mechanism distinct from that in *Arabidopsis*. FG-*siren* loci represent a small set of high-output 24-nt siRNA-producing regions that dominate the female reproductive siRNA landscape in rice, undergo progressive CHH methylation during FG development, and are distinct from endosperm *siren* loci. Together, these findings reveal an intrinsic motif-binding capacity of a plant CLSY protein and a distinct CLSY-mediated RdDM targeting mode independent of both histone methylation and transcription factors.

RdDM-driven CHH methylation ensures rice reproduction^20–26^. Our study shows that OsRDR2 generates the majority of 24-nt siRNAs from FG-*siren* loci in female reproductive tissues and panicles, although FG-*siren*-overlapping DMRs account for only a small fraction of all *osrdr2* hypo-DMRs in these organs (Fig. 3 and Supplementary Figs. 9,10). Notably, CHH methylation is strongly reduced across *osrdr2* hypo-DMRs, despite the absence of clear, autonomous, OsRDR2-dependent siRNA production from non-overlapping DMRs in these tissues (Fig. 3c–f). Given the dominant contribution of FG-*siren* loci to 24-nt siRNA production, we cannot rule out the possibility that FG-*siren*-derived siRNAs regulate CHH methylation at other hypo-DMRs in *trans*. However, in previously reported cases of *trans* RdDM, the number of trans-regulated targets was comparable to that of siRNA source loci^40,61^, suggesting that potential *trans* RdDM alone is unlikely to fully explain the large disparity in number between FG-*siren* loci and *osrdr2* hypo-DMRs. Alternatively, the large number of non-overlapping hypo-DMRs may reflect the cumulative consequence of constitutive OsRDR2 loss throughout the developmental trajectory leading to reproductive organs. In this scenario, some CHH methylation defects detected in pistils may originate from loss of RdDM activity at earlier developmental stages across the vegetative-to-reproductive transition, whereas only pistil-active RdDM targets are expected to show autonomous siRNA production in pistils. Thus, FG-*siren* loci may represent high-output reproductive siRNA sources, whereas the broader *osrdr2* methylation defects likely reflect both local and developmentally accumulated consequences of impaired RdDM.

We observed distinct siRNA pools in female reproductive tissues and developing endosperm (Fig. 2f and Supplementary Fig. 7), as previously reported^32^, indicating distinct RdDM regulation between rice endosperm and female reproductive tissues. Brassicaceae ovules and endosperms share *siren*-derived siRNAs to a considerable extent, presumably through non-cell-autonomous movement of maternal siRNAs^16,30^. Nonetheless, the presence of maternal sporophytic siRNAs in rice endosperm (Fig. 2g and Supplementary Fig. 8) suggests that similar maternal siRNA transport may occur in rice endosperm, albeit to a lesser extent. This may reflect a true biological difference in siRNA movement between monocots and dicots; alternatively, transported siRNAs may be diluted during the rapid expansion of rice endosperm, which involves endoreduplication, cellularization, and storage product accumulation^62,63^. However, a partial contribution from trace seed coat contamination cannot be completely excluded. This separation between female reproductive and endosperm RdDM is further reflected in *osclsy* mutant analyses and the genetic requirements for siRNA accumulation. Despite the distinct regulation of FG-*siren* and endosperm *siren* loci (Fig. 2f and Supplementary Fig. 7), siRNA expression at FG-*siren* loci was reduced in *osclsy4-kd* endosperm rather than in *osclsy3-kd* endosperm (Supplementary Fig. 12). However, OsCLSY4 did not bind the FRE motif in vitro (Supplementary Fig. 18). These results suggest that RdDM in endosperm is regulated through a molecular mechanism distinct from the OsCLSY3-FRE module active in female reproductive tissues. Given the redundant but unique roles of *OsCLSY4* and *OsCLSY3* during endosperm development^22^, future studies will be needed to determine how endosperm *siren* loci are specified in rice and whether similar separation between female reproductive and endosperm siRNA pools occurs broadly across monocots.

Tissue-specific divergence may also extend to male reproductive tissues. In Arabidopsis, CLSY3 promotes HyperTE-derived 24-nt siRNA production in tapetal nurse cells, from which these siRNAs move to meiocytes and guide methylation at TEs and genes^61^. Whether similar intercellular communication occurs in rice anthers remains to be explored. However, weak FG-*siren*-derived siRNA expression in rice stamens and marginal *OsCLSY3* expression in male reproductive tissues do not support a major role for OsCLSY3 in rice male reproductive RdDM (Fig. 2a and Supplementary Figs. 9a,18b). Instead, premeiotic and meiotic phasiRNAs may largely shape the DNA methylation landscape in rice anthers, as suggested by their predominance during male reproduction in grasses^64^ and contribution to CHH methylation in maize^65^.

The intrinsic FRE-binding activity of OsCLSY3 raises the question of how a CLSY chromatin remodeler can recognize a specific DNA motif. CLSY proteins are SNF2-related chromatin remodeling factors containing conserved SNF2_N and Helicase_C domains, which couple ATP hydrolysis to chromatin remodeling activity^66,67^. Because SNF2-family remodelers interact with DNA as part of their remodeling activity, their DNA engagement is generally considered sequence-independent^66^, and sequence-specific motif recognition has not been established for plant CLSY proteins. However, preferential interactions with GC-rich or structurally distinctive DNA have been reported for other SNF2 proteins, including human ATRX, which targets GC-rich tandem repeats and binds G-quadruplex structures in vitro, and maize ZmDDM1, which autonomously recognizes a GC-rich motif during RdDM-mediated maintenance of CHH islands^68,69^. The FRE motif has distinctive sequence features that may contribute to OsCLSY3 recognition. It is palindromic and GC-rich, with conserved terminal GC-rich cores and an internal A/T-rich region, and the association between FRE and FG-*siren* loci is sensitive to both features (Supplementary Fig. 13). This architecture may influence DNA shape or local structural properties, such as hairpin- or cruciform-like conformations, rather than canonical G-quadruplex formation, which requires repeated G-tracts^70,71^. Together with the lack of FRE binding by OsCLSY4 (Supplementary Fig. 18), these features suggest that FRE recognition is unlikely to be a general property of CLSY proteins but may instead reflect a specialized activity of OsCLSY3. Future structural and biochemical studies, together with motif perturbation, will be important for determining whether FRE sequence composition directly instructs OsCLSY3 recruitment and RdDM activity.

RdDM has different impacts on reproductive development across plant species^13–26^. In rice, RdDM-mediated CHH methylation is crucial for reproductive success^20–25^, and our results show that FG-*siren* loci serve as major RdDM targets during reproductive development (**Fig. 3**), suggesting that CHH methylation in these regions may play a critical role in rice reproduction. In maize, mutation of the RDR2 ortholog *Mop1* (*Mediator of paramutation 1*) affects recombination landscapes by modulating euchromatic TEs near genes^18,19^. In Brassicaceae, *B. rapa* shares high *CLSY3* expression in ovules and enrichment of AT-rich TE-associated motifs at *siren* loci with *Arabidopsis*^40^. However, ovule *siren* loci in *Arabidopsis* and *B. rapa* are largely found in non-conserved genomic regions^30^, indicating a shared mode of production but divergent genomic distribution of ovule *siren* siRNAs between these species. Such differences in genomic distribution may contribute to their different reproductive dependence on RdDM, with maternal RdDM being required for *B. rapa* seed development^14^, whereas RdDM is largely dispensable for *Arabidopsis* reproduction^72^. Our results extend this idea to rice, in which female-preferential RdDM differs from Arabidopsis not only in genomic distribution but also in motif composition and CLSY3-mediated motif recognition. Rice FG-*siren* loci are preferentially distributed at genic TEs on euchromatic chromosomal arms (Fig. 1), consistent with reports that OsDRM2 is required for CHH methylation of gene-associated TEs^51^ and that CHH islands occur near genes in maize and rice but are largely absent in *Arabidopsis*^73,74^. FG-*siren* loci are enriched for the GC-rich FRE motif (Fig. 4) and associated with direct interaction between OsCLSY3 and FRE-containing DNA (Fig. 5). In contrast, Arabidopsis reproductive RdDM is associated with AT-rich motifs and requires REM transcription factors and GDE1 for CLSY3/4-mediated targeting^38,39^. Thus, although in principle CLSY3-mediated, DNA motif-associated, female-enriched RdDM appears to be conserved, the molecular basis of motif recognition and the resulting genomic distribution of RdDM targets may differ across lineages. Given the proposed Poaceae-specific duplication within *CLSY3/4* clades^60^, these findings further suggest that OsCLSY3 may have acquired an intrinsic sequence-specific motif-binding capability through neofunctionalization, providing a possible mechanism for the distinct genomic targeting and reproductive impact of RdDM in rice.

## Methods

### Plant materials, growth conditions, and sample collection

*Oryza sativa* L. ssp. *japonica* cv. Nipponbare was used in this study. Dehulled seeds were surface-sterilized with 70% ethanol for 1 min followed by 1.5% sodium hypochlorite for 30 min and germinated on half-strength Murashige and Skoog (MS) medium supplemented with 3.0% (w/v) sucrose and 0.3% (w/v) Gelrite. Seedlings were incubated in an AR-36 growth chamber (Percival Scientific) under continuous light (4000 lux) at 28 °C. After 10 days, seedlings were transplanted into a 5:2 mixture of Pro-Mix HP Mycorrhizae (Premier Tech Horticulture) and Sun Gro® Peat Moss Grower Grade White (Sun Gro Horticulture). Plants were grown in a greenhouse in Gainesville, Florida, USA (29.65° N), under natural light conditions, with an approximately 14 h photoperiod and day/night temperatures of 28 °C/25 °C.

Developmental stages of ovaries were assigned based on pistil length, as previously described^41^. Briefly, immature rice spikelets were collected, and pistils were extracted using fine forceps. Pistil length was measured using a Leica MSV269 microscope, and samples were classified into six stages: megaspore mother cell stage (MMC; <1.1 mm), meiosis stage (MEI; 1.1–1.4 mm), early mitosis stage (MIT1; 1.4–1.7 mm), late mitosis stage (MIT2; 1.7–2.0 mm), early mature stage (MAT1; 2.0–2.4 mm), and late mature stage (MAT2; >2.4 mm). Pistils were separated into ovary and stigma/style fractions using a razor blade, and only ovaries were collected and frozen in liquid nitrogen. Flag leaves during the reproductive phase were collected and frozen in liquid nitrogen and used as vegetative controls.

### sRNA-seq library preparation, sequencing, and mapping

For small RNA sequencing (sRNA-seq), at least 50 ovaries were pooled for each library.

Total RNA was extracted from ovaries and flag leaves using the RNeasy Plant Mini Kit (Qiagen), coupled with on-column DNase I treatment using RNase-Free DNase Set (Qiagen). Libraries were prepared using the NEBNext® Multiplex Small RNA Library Prep Set for Illumina® (NEB). Library products were purified using an 8% polyacrylamide gel, and fragments of 140–160 bp, corresponding to sRNA inserts plus 3’ and 5’ adapters, were excised. Sequencing was performed on a NovaSeq 6000 platform.

Single-end 50-bp reads were trimmed and filtered using TrimGalore (v0.6.10) with default parameters except that the minimum length was set to 10 bp (--length 10). Cleaned reads were mapped to the rice Nipponbare reference genome IRGSP-1.0^76^ using Bowtie (v1.2.3)^77^ with no mismatches allowed (-v 0). Uniquely mapped reads were retained (-m 1). Reads mapped to rRNA, tRNA, snRNA, snoRNA (based on RNAcentral v22^78^), and miRNA (based on miRBase v22^79^) were removed using bedtools (v2.30.0)^80^ intersect, and the remaining reads were considered siRNA reads. Uniquely mapped 24-nt and 21–22-nt siRNA reads were used for further analyses. BigWig tracks were generated using bamCoverage from deepTools (v3.5.2) package^81^ with CPM normalization (--normalizeUsing CPM).

Publicly available sRNA-seq datasets from vegetative and reproductive tissues of different rice varieties and RdDM-related mutants were retrieved from public repositories, including NCBI GEO and BioProject, under the following accessions: GSE20748, GSE22763, GSE26405, GSE44989, GSE50778, GSE81436, GSE90479, GSE107904, GSE108527, GSE112259, GSE130122, GSE130168, GSE131319, GSE152155, GSE154501, GSE158711, GSE180457, GSE215857, GSE229961, GSE290279, PRJNA533115, and PRJDB7574. Raw reads were processed using the same pipeline described above, with dataset-specific adapter trimming where applicable. All sRNA-seq datasets used in this study and their mapping statistics are summarized in Supplementary Data 1.

### EM-seq library preparation, sequencing, and mapping

For enzymatic methylation sequencing (EM-seq), at least 35 ovaries were pooled for each library. Genomic DNA was extracted from ovaries and flag leaves using the DNeasy Plant Mini Kit (Qiagen). EM-seq libraries were constructed using the NEBNext® Enzymatic Methyl-seq Kit (New England Biolabs) according to the manufacturer’s instructions. Sequencing was performed on the Illumina NovaSeq 6000 platform.

Raw reads were quality-checked using FastQC (v0.11.7; https://www.bioinformatics.babraham.ac.uk/projects/fastqc/). Paired-end reads were trimmed with Trimmomatic (v0.39)^82^ with the following parameters: LEADING:3, TRAILING:3, SLIDINGWINDOW:4:20, MINLEN:36. Clean reads were aligned to the reference genome IRGSP-1.0^76^ using Bismark v0.22.3 (-I 50 -N 1)^83^ with bowtie2 as the aligner^84^. PCR duplicates were removed using deduplicate_bismark. Methylation calling was performed using bismark_methylation_extractor, and bismark2bedGraph, and coverage2cytosine commands. The bedGraph files of each sample for each sequence context (CG, CHG, or CHH) were generated by calculating methylation rates for each cytosine supported by at least four mapped reads. BigWig tracks were subsequently generated using bedGraphToBigWig command from UCSC tools^85^.

Available BS-seq datasets from vegetative and reproductive tissues of different rice varieties and RdDM-related mutants were retrieved from public repositories, including NCBI GEO and BioProject, under the following accessions: GSE22591, GSE130122, GSE81436, GSE89789, GSE112259, GSE130168, GSE152155, GSE158711, PRJNA533115, and PRJDB7574. Raw reads were processed using the same pipeline described above, with dataset-specific read-layout adjustments where applicable. All EM-seq and BS-seq datasets used in this study and their mapping statistics are summarized in Supplementary Data 2.

### 24-nt siRNA and DNA methylation data analyses and visualization

#### 24-nt siRNA cluster calling

BAM files of 24-nt siRNA reads from replicates were merged using samtools (v.1.20)^86^ merge and subsequently converted into BED files using bedtools^80^ bamtobed. Clusters were identified using bedtools^80^ merge (-d 200) and read counts per cluster were counted using bedtools^80^ coverage. Clusters with reads per million (RPM) > 10 of 24-nt siRNAs were retained for further analyses. Cumulative sums of 24-nt siRNA levels were calculated by sorting clusters in descending order based on their expression levels. A cumulative sum plot of 24-nt siRNAs for each sample was generated using the ggplot2 package^87^ in R. 24-nt siRNA abundant loci were identified at the knee point using the Kneedle algorithm^88^ and are provided in Supplementary Data 4. Common 24-nt siRNA abundant loci across six stages of developing ovaries were determined using bedtools^80^ multiinter. Scaled Venn diagrams showing the overlaps among 24-nt abundant loci across six stages of developing ovaries were generated using the eulerr package^89^ in R. A final set of FG-*siren* loci (137 loci) was generated using bedtools^80^ merge (-d 1000) and are provided in Supplementary Data 5.

The overlap between FG-*siren* loci and genomic features, including genes, transposons, pericentromeric heterochromatin were obtained using bedtools^80^ intersect. The overrepresented transposon families were examined by comparing the observed overlap with the expected overlap based on their genomic occupancy. Statistical significance was determined using the Fisher’s exact test.

#### 24-nt siRNA expression and DNA methylation

For boxplots, the expression levels of 24-nt siRNAs for different samples at the indicated regions were obtained using bedtools^80^ multicov. Corresponding DNA methylation levels were calculated based on cytosines supported by at least 10 mapped reads. RPM-normalized siRNA levels and DNA methylation levels were visualized as boxplots using the ggplot2 package^87^ in R. Proportion of mapped 24-nt siRNA reads to total reads was visualized as bar plots using the ggplot2 package^87^ in R.

Pairwise relationships among 24-nt siRNA expression levels (log_2_(1+RPM)) and DNA methylation levels (%) across developing ovaries and flag leaves were visualized as scatter plots using the ggplot2 package^87^ in R. The corresponding Pearson correlation coefficients were calculated, and statistical significance was determined using Benjamini-Hochberg (BH)-adjusted false discovery rates (FDR).

#### Overlap among 24-nt siRNA clusters and DMRs

Genomic coordinates for the 24-nt siRNA clusters and DMRs were obtained from previous studies^20–23,27,28,42,53^ and are listed in **Supplementary Data 7–10**. Among these, endosperm imprinted sRNA loci were defined using MSU v6 genome assembly^27^; their genome coordinates were therefore converted to IRGSP-1.0 by obtaining sequences using bedtools^80^ getfasta, mapping the sequences to the IRGSP-1.0 genome^76^ using Bowtie^77^, and converting the alignments to genome coordinates using bedtools^80^ bamtobed.

Scaled Venn diagrams showing the relationship among 24-nt siRNA clusters and CHH hyper- or hypo-DMRs were generated using the eulerr package^89^ in R. 24-nt or 21–22-nt siRNA reads mapped to each intersection category were obtained using bedtools^80^ intersect and their proportions were visualized using pie charts.

#### sRNA read composition

BAM files of 24-nt and 21–22-nt siRNA reads were compared with heterochromatic siRNAs (hc-siRNAs) and 21-nt or 24-nt phased secondary siRNAs (phasiRNAs) (based on sRNAanno^90^) or different families of transposable elements using bedtools^80^ intersect.

### Motif discovery and data analysis

DNA motifs enriched at FG-*siren* loci were identified using STREME^91^ under MEME Suite v5.4.1^92^. Genome-wide occurrences of the motifs were assessed using FIMO^93^ with the P-value under 0.0001 (--thresh 1.0E-4) under MEME Suite^92^. The motif occurrences at FG-*siren* loci were subsequently assessed using bedtools^80^ intersect. The motif positions on FG-*siren* loci were calculated in 20 proportional bins. The number of mismatches in the FRE core sequences and GC-content in the internal sequences were assessed using a custom script. Their effects on the association between FRE motif and FG-*siren* loci were evaluated using the Cochran-Armitage trend test. The motif distribution and occurrence in FG-*siren* loci were visualized using the ggplot2 package^87^ in R.

Heatmaps showing the association between a DNA motif and 24-nt siRNA and DNA methylation levels at FG-*siren* loci were generated using deepTools^81^. The representative motifs per locus were determined as the motif with the highest score, where the motif score indicates the log-likelihood similarity to the consensus motif sequence. Matrix files were generated using computeMatrix (reference-point -a 5000 -b 5000 --binSize 50 --sortRegions keep), and heatmaps were drawn using plotHeatmap.

### Genome-wide distribution of FG-*siren* loci and FRE motifs

The genome-wide distribution of genomic features, FG-*siren* and endosperm *siren* loci, 24-nt siRNA levels of Nipponbare panicles, ovary (MAT2 stage), and endosperm were visualized using the ggplot2 package^87^ in R. For each 100-kb genomic bin, the percentage of the bin covered by genes, transposons, FG-*siren* loci, and endosperm *siren* loci was calculated. The maximum FRE motif score within each bin was shown. 24-nt siRNA levels per bin were determined using bedtools^80^ multicov and log_2_(1+RPM)-normalized values were shown.

### mRNA-seq library preparation, sequencing, and mapping

For messenger RNA sequencing (mRNA-seq), at least 50 ovaries were pooled for each library. Total RNA was extracted from ovaries and flag leaves using the RNeasy Plant Mini Kit (Qiagen), coupled with on-column DNase I treatment using RNase-Free DNase Set (Qiagen). mRNA was selected using the NEBNext® Poly(A) mRNA Magnetic Isolation Module (NEB), and libraries were prepared with the NEBNext® Ultra™ II RNA Library Prep Kit for Illumina® (NEB). Sequencing was performed on the NovaSeq 6000 platform (Illumina).

The quality of the raw reads was assessed using FastQC. Paired-end reads were cleaned using Trimmomatic (LEADING:3, TRAILING:3, SLIDINGWINDOW:4:20, MINLEN:36)^82^.

Trimmed reads were then aligned to the rice genome IRGSP-1.0^76^ using HISAT2 (v2.2.1)^94^. Gene-level read counts were calculated based on RGAP 7 annotation^95^ using HTSeq-count (v0.11.2)^96^ with the “union” resolution mode, excluding reads mapped to more than one gene (-- mode union --nonunique none). Transcripts per million (TPM) values were calculated using a custom script and used for downstream analyses.

Public mRNA-seq datasets from vegetative and reproductive tissues of different rice varieties were retrieved from repositories, including NCBI GEO and BioProject, under the following accessions: GSE50778, GSE107903, GSE112259, GSE130122, GSE130168, GSE131319, GSE152155, GSE154501, GSE180457, GSE229961, and PRJDB7574. Raw reads were processed using the same pipeline described above. All mRNA-seq datasets used in this study and their mapping statistics are summarized in **Supplementary Data 3**.

### Correlation between gene expression and 24-nt siRNA levels

Pearson correlation coefficients were calculated for each expressed gene between its transcript levels and 24-nt siRNA levels at FG-*siren* loci across 22 vegetative and reproductive tissues. Genes were sorted in ascending order based on their correlation coefficients and visualized using the ggplot2 package^87^ in R.

### Transcript levels of OsCLSY genes

TPM-normalized transcript levels of *OsCLSY1* (LOC_Os07g49210), *OsCLSY4* (LOC_Os05g32610), and *OsCLSY3* (LOC_Os02g43460) across vegetative and reproductive tissues were visualized as bar plots using the ggplot2 package^87^ in R.

### ChIP-seq mapping and data analysis

ChIP-seq of OsCLSY3 tagged with C-terminal epitopes (GFP and 2xMYC) in rice panicles^21^ were analyzed. Raw reads were obtained from NCBI GEO database (GSE229961), trimmed using Cutadapt v5.0^97^, and aligned to IRGSP-1.0 genome^76^ using Bowtie2 v2.5.4^84^ with --very-sensitive-local -N 1 parameters. The PCR duplicates were removed using Picard v2.16.0 (http://broadinstitute.github.io/picard/). TagDirectories from ChIP-seq data were generated using the HOMER^98^ makeTagDirectory script. OsCLSY3 ChIP peaks were identified using findPeaks function from HOMER^98^ (-style factor -L 3 -F 3 -center -region). Peaks from two replicates were integrated using the HOMER^98^ mergePeaks script (-d given) and filtered using rice greenscreen regions obtained from Klasfeld, Roulé, and Wagner (2022)^99^ to remove artifactual signals.

Detailed information on ChIP peaks is provided in Supplementary Data 14.

Intersections between OsCLSY3 ChIP peaks and FG-*siren* loci were determined using bedtools^80^ intersect and visualized as scaled Venn diagram using the eulerr package^89^ in R. Detailed information of their intersection is provided in Supplementary Data 16. HOMER^98^ findMotifsGenome.pl script (-size 200 -len 12,16,20 -p 10 -mis 4 -bits) was used to examine FRE enrichment and de novo identify DNA motifs found in OsCLSY3 peaks. DNA motifs enriched in ChIP peaks are provided in Supplementary Data 15.

OsCLSY3 ChIP enrichment at the identified peaks and FG-*siren* loci were visualized using deepTools^81^. Briefly, the BAM files were converted to bigWig format using the bamCoverage command (-bs 25 --normalizeUsing RPGC), and ChIP signals were compared with Input controls using the bigwigCompare command (--operation log2). Resulting bigWig files were used to generate data matrices using the computeMatrix script (reference-point -- referencePoint center -a 5000 -b 5000 -bs 200), and visualized using plotHeatmap and plotProfile.

### Prediction of protein-DNA interaction using AlphaFold3

The interaction between OsCLSY3 and DNA probes containing or lacking the FRE motif was predicted using AlphaFold3 (AF3)^59^ through web server (https://alphafoldserver.com/). Top-ranked prediction models for each DNA probe were visualized using PyMOL v3.0.5. A detailed list of probes used for AF3 predictions is provided in Supplementary Data 17.

### EMSA

Coding regions corresponding to OsCLSY4 (amino acids 850-1,445) and OsCLSY3 (amino acids 870-1,439), containing SNF2-related (PF00176) and Helicase domains (PF00271), were synthesized and cloned into pET-30a (+). Recombinant plasmids were transformed into *E. coli* strain BL21 (DE3), and protein expression was induced with 0.5 mM IPTG. Cells were lysed, inclusion bodies were washed, and proteins were purified using Ni-NTA resin (GenScript). Proteins were subsequently refolded in a buffer containing 50 mM Tris-HCl, 150 mM NaCl, 1 M L-Arginine, 10 % glycerol, pH 8.0.

For EMSA, 2 µg of recombinant proteins were incubated with 4 pmol of biotin-labeled probes with or without cold probes in EMSA/Gel-shift binding buffer (Beyotime) supplemented with 5 µM ATP and 5 µM MgCl_2_ at 25 °C for 30 minutes. Reaction samples were separated using 6% native polyacrylamide gel electrophoresis (PAGE). After transferring to the PVDF membrane, biotin-labeled probes were detected using HRP-conjugated streptavidin (Proteintech) and an ECL detection kit (Vigorous Biotech). The probes used for EMSA are listed in Supplementary Data 17.

### Statistics and reproducibility

Overrepresentation of transposon families at FG-*siren* loci was assessed using the Fisher’s exact test. Pairwise correlation of 24-nt siRNA and DNA methylation levels at FG-*siren* loci was determined based on the Pearson correlation coefficient and BH-adjusted FDR. For all boxplots, the boxes show the interquartile ranges (IQR), with the black lines representing the median, and the whiskers extend no longer than 1.5 times the IQR.

### Data availability

Illumina sequencing of EM-seq, mRNA-seq, and sRNA-seq datasets generated in this study were deposited in the NCBI Gene Expression Omnibus (GEO) and are accessible via accession numbers GSE282528, GSE282529, and GSE282531, respectively. Published datasets used in this study include the followings: GSE20748 (Ref. ^49^), GSE22591 (Ref. ^54^), GSE22763 (Ref. ^46^), GSE26405 (Ref. ^47^), GSE44989 (Ref. ^27^), GSE50778 (Ref. ^50^), GSE81436 (Ref. ^51^), GSE89789 (Ref. ^55^), GSE90479 (Ref. ^45^), GSE107903, GSE107904 (Ref. ^48^), GSE108527 (Ref. ^52^), GSE112259, GSE130168, GSE152155 (Ref. ^20^), GSE130122 (Ref. ^28^), GSE131319 (Ref. ^53^), GSE154501 (Ref. ^42^), GSE158711 (Ref. ^24^), GSE180457 (Ref. ^26^), GSE215857 (Ref. ^23^), GSE229961 (Ref. ^21^), GSE290279 (Ref. ^22^), PRJNA533115 (Ref. ^31,32^), and PRJDB7574 (Ref.^44^). All custom scripts are available from the corresponding author upon request. Source data are provided with this paper.

## Acknowledgements

We gratefully acknowledge Evert Varela for providing technical support with the isolation of rice samples during the initial stages of this project. This work was supported by the Hatch project FLA-ENH-006627 from the United States Department of Agriculture and the Competitive Seed Grant Research Initiative (Grant No. 00129910) from the College of Agricultural and Life Sciences at the University of Florida to K.B.

## Author contributions

K.B. conceptualized the study. K.B. and T.K. designed the experiments. T.K. performed bioinformatic analyses and experiments. M.Z. provided bioinformatic support. M.F.R.R. provided computational infrastructure. K.B. and T.K. interpreted the data and wrote the manuscript. All authors read and approved the manuscript.

