## Supplementary Figures for "Intrinsic DNA motif-binding capability of OsCLSY3 guides female-preferential RNA-directed DNA methylation"

**Supplementary Figs. 1–18**

**Supplementary Note 1**
**Supplementary Note Figs. 1–2**

**Supplementary Data 1–17** (available as **Additional Files**)

**Supplementary References**

### Supplementary Figures

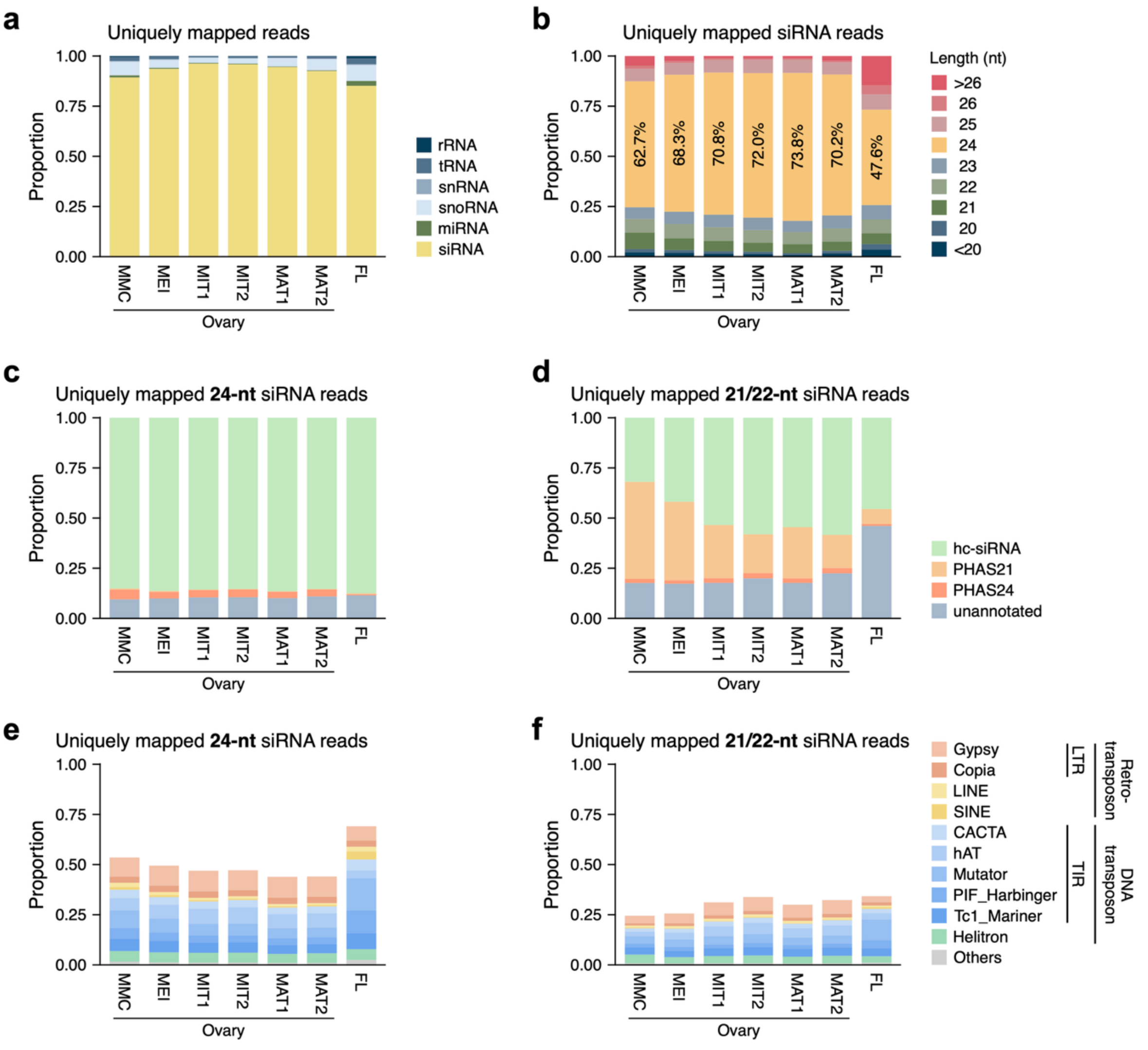


**Supplementary Fig. 1. sRNA composition across tissues. a,** Proportion of structural RNAs (rRNAs, tRNAs, snRNAs, and snoRNAs), miRNAs, and siRNAs among uniquely mapped sRNA reads. **b,** Length distribution of uniquely mapped siRNA reads. **c,d,** Proportion of hc-siRNAs and phasiRNAs within 24-nt siRNAs (**c**) and 21–22-nt siRNAs (**d**). **e,f,** Proportion of 24-nt (**e**) and 21–22-nt (**f**) siRNA reads mapping to transposons. sRNA-seq datasets are listed in Supplementary Data 1.

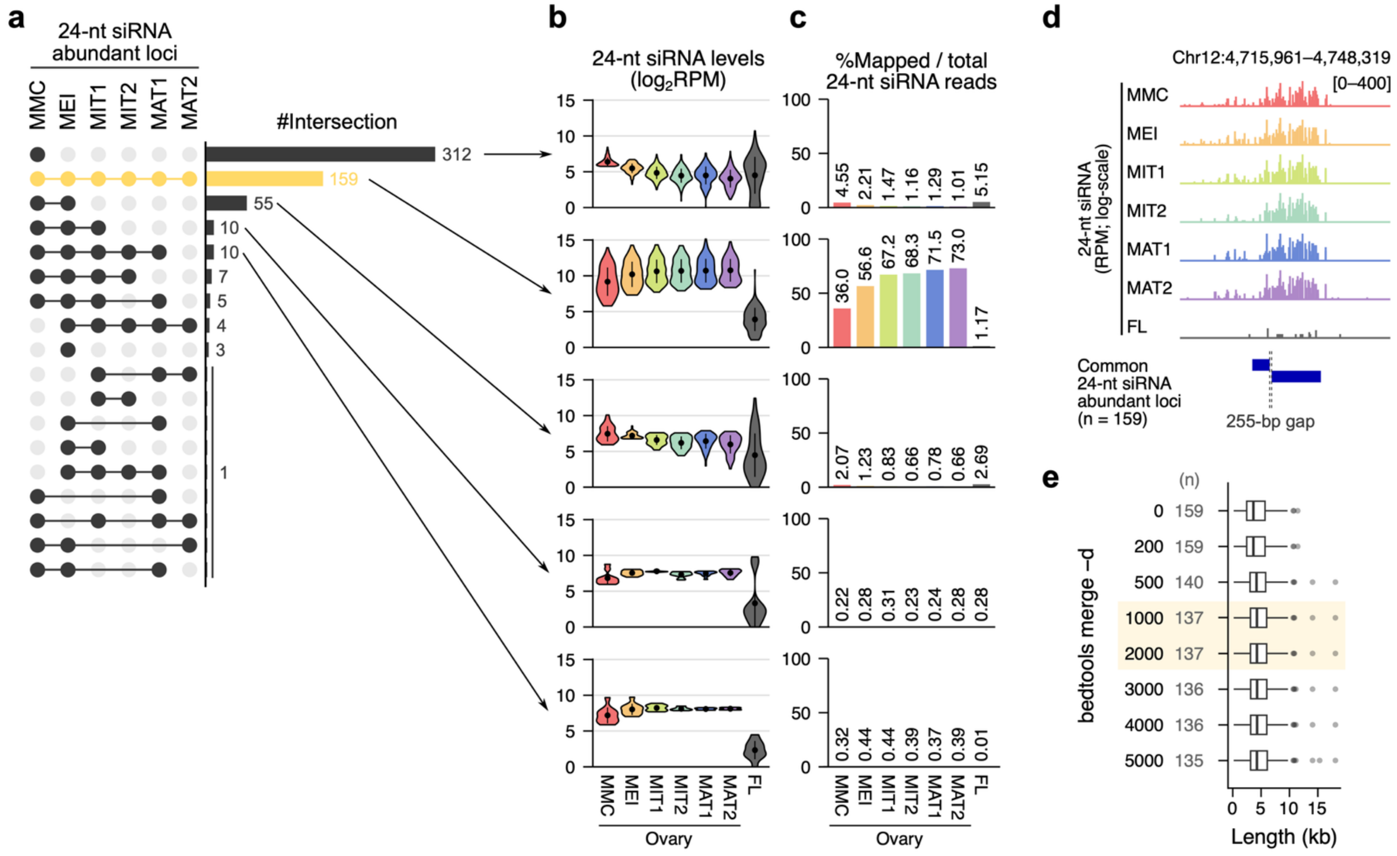


**Supplementary Fig. 2. Definition of FG-*siren* loci. a,** Relationship among 24-nt siRNA abundant loci in ovaries across developmental stages. Common 24-nt siRNA abundant loci across all developmental stages are highlighted. **b,c,** 24-nt siRNA levels at each locus (**b**) and the proportion of 24-nt siRNA reads mapped to all loci (**c**) across the intersection developmental categories containing ≥10 regions. **d,** Length distribution of FG-*siren* loci obtained at each merge-gap threshold; the selected threshold and resulting final set of FG-*siren* loci (137 loci) are highlighted in yellow. For boxplots, center lines indicate medians, boxes indicate interquartile ranges, and whiskers extend no further than 1.5 times the interquartile range; points indicate outliers. sRNA-seq datasets are listed in Supplementary Data 1. Genomic coordinates of FG-*siren* loci are listed in Supplementary Data 5.

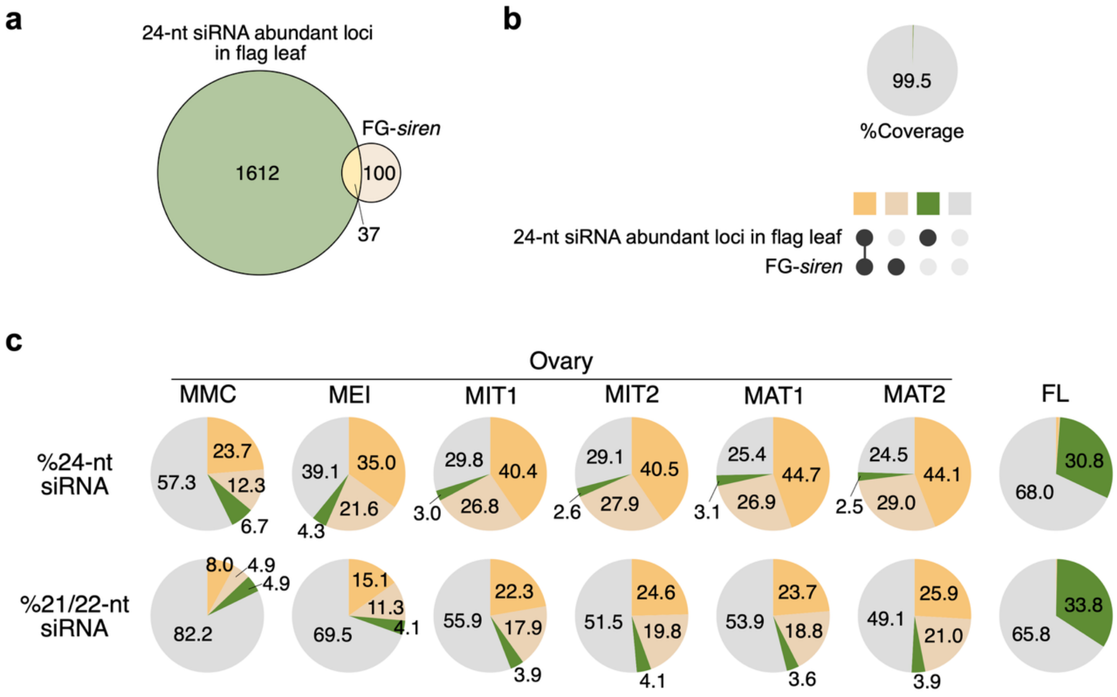


**Supplementary Fig. 3. Comparison of 24-nt siRNA abundant loci in ovaries and flag leaves. a,** Relationship between 24-nt siRNA abundant loci in flag leaves and FG-*siren* loci. **b,** Genome coverage of each intersection category. **c,** Relative abundance of 24-nt and 21–22-nt siRNAs expressed in each intersection category. sRNA-seq datasets are listed in Supplementary Data 1. Genome coordinates for 24-nt siRNA abundant loci in flag leaves and comparisons with FG-*siren* loci are listed in Supplementary Data 6.

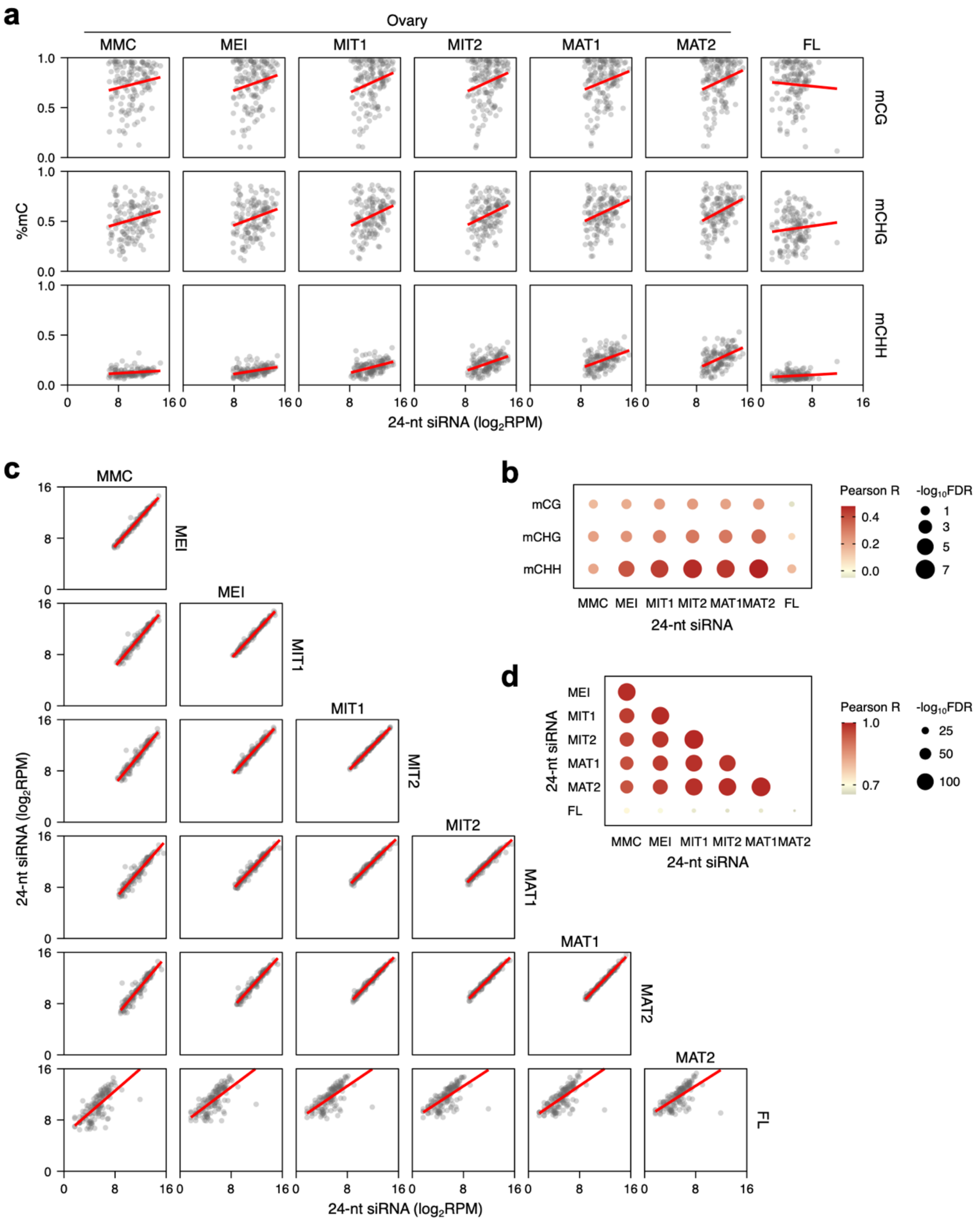


**Supplementary Fig. 4. Comparison of 24-nt siRNA accumulation and CHH methylation at FG-*siren* loci in ovaries and flag leaves. a,b,** Correlations between 24-nt siRNA and CHH methylation levels at FG-*siren* loci in ovaries and flag leaves and the corresponding Pearson correlation coefficient (*R*) and Benjamini-Hochberg (BH)-adjusted false discovery rates (*q*) (**b**). **c,d,** Pairwise correlations of 24-nt siRNA levels across tissues (**c**) and the corresponding *R* and *q* values (**d**). For **a**,**c**, red lines represent simple linear regression fits. sRNA-seq datasets are listed in Supplementary Data 1.

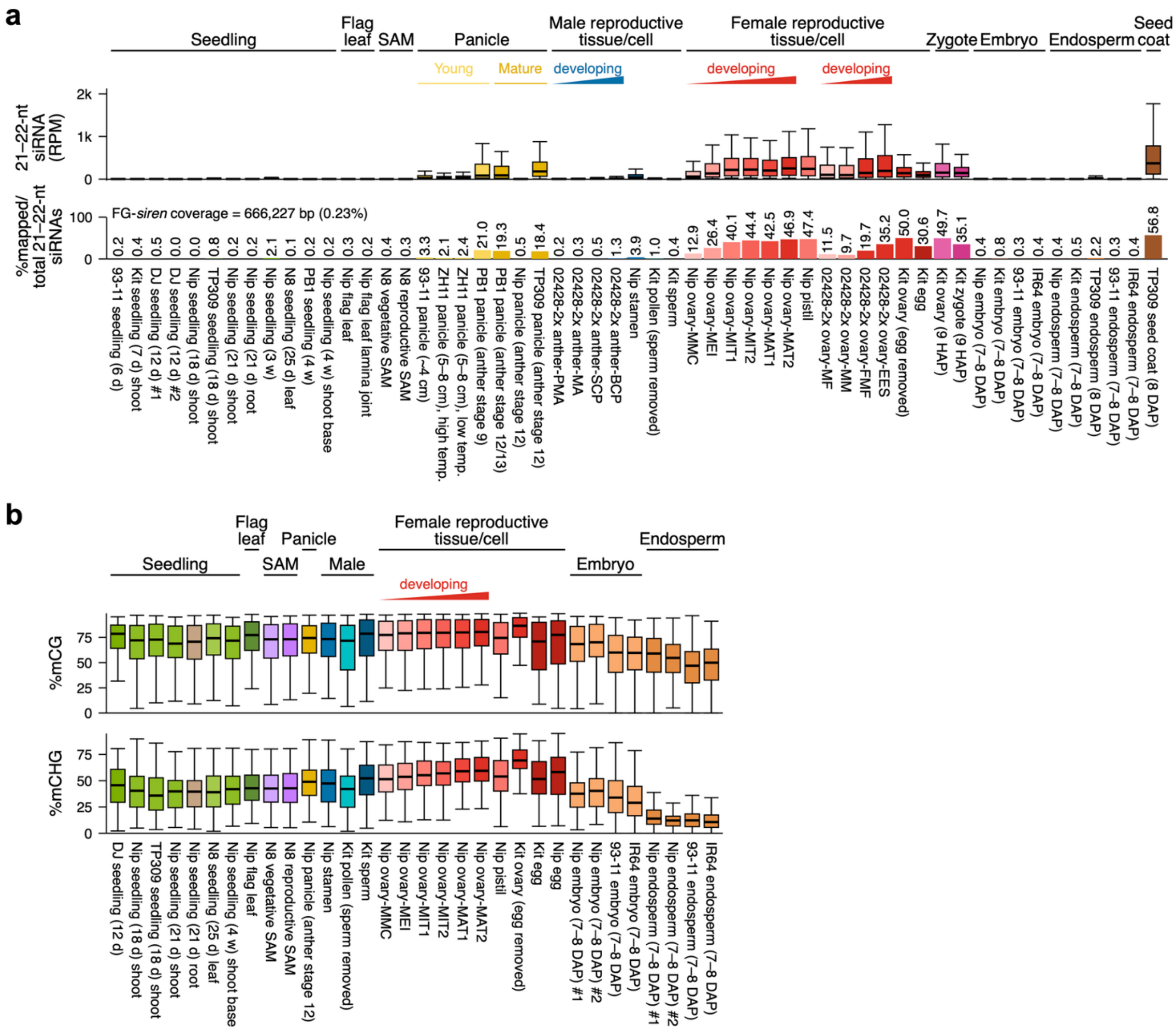


**Supplementary Fig. 5. Comparison of siRNA expression and DNA methylation levels at FG-*siren* loci across tissues and genotypes. a,b,** 21–22-nt siRNA expression (**a**) and CG and CHG methylation levels (**b**) at FG-*siren* loci across vegetative and reproductive tissues from different rice varieties. In the lower panel of **a**, the relative abundance of 21–22-nt siRNAs derived from FG-*siren* loci is shown. sRNA-seq and BS-seq datasets obtained from public repositories[1–23] and are listed in Supplementary Data 1 and 2, respectively. Abbreviations for rice cultivars: Nip, Nipponbare; Kit, Kitaake; TP309, Taipei 309; ZH11, Zhonghua 11; N8, Norin 8; DJ, Dongjin; PB1, Pusa Basmati 1.

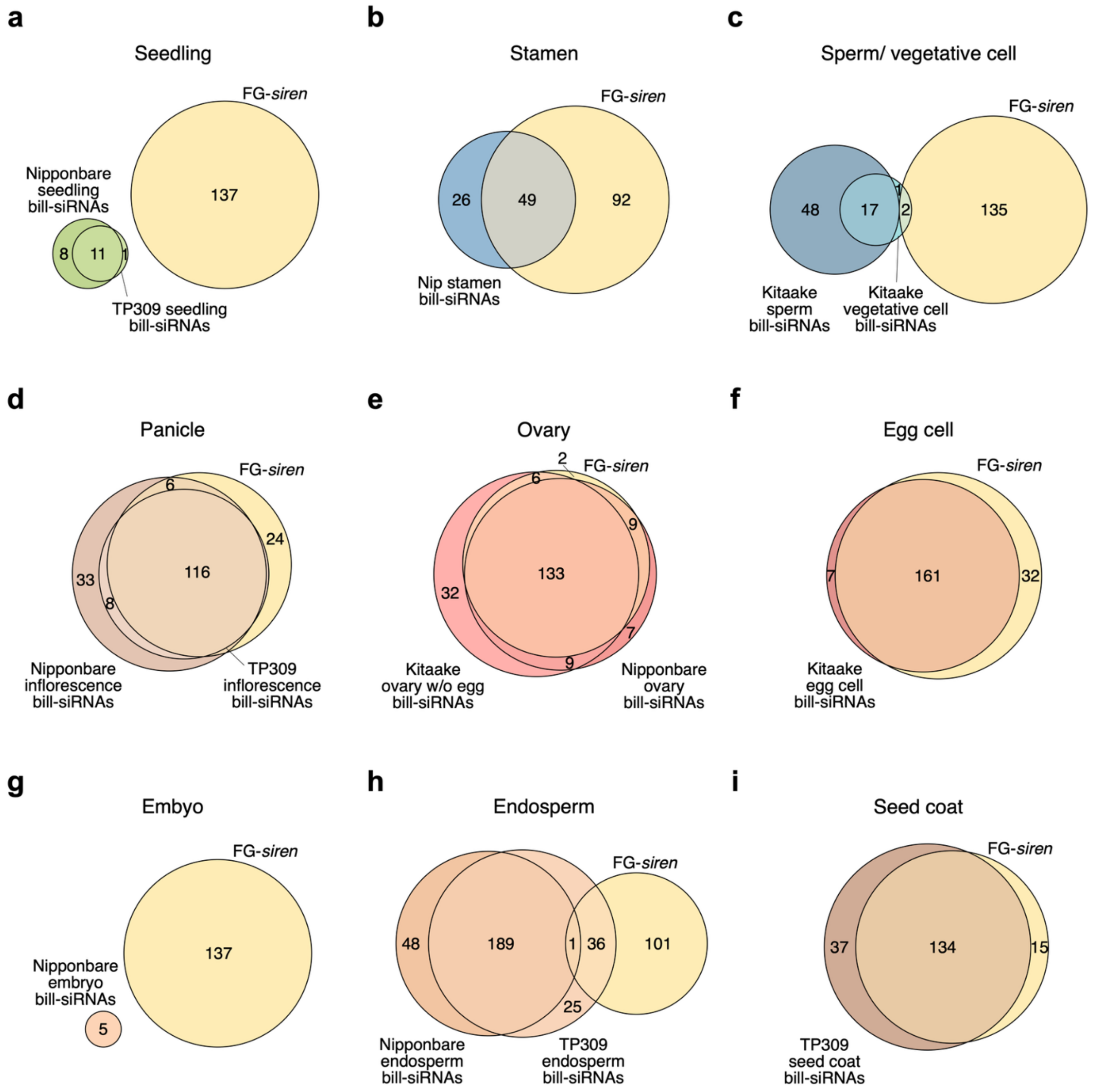


**Supplementary Fig. 6. Comparison of FG-*siren* loci with highly expressed siRNA clusters across vegetative and reproductive tissues. a–i,** Overlap between FG-*siren* loci and billionaire siRNA clusters (bill-siRNAs) identified from seedling (**a**), stamen (**b**), sperm and vegetative cell (**c**), panicle (**d**), ovary (**e**), egg cell (**f**), embryo (**g**), endosperm (**h**), and seed coat (**i**). Genome coordinates for bill-siRNAs were obtained from a previous study[19], and comparisons with FG-*siren* loci are listed in Supplementary Data 7.

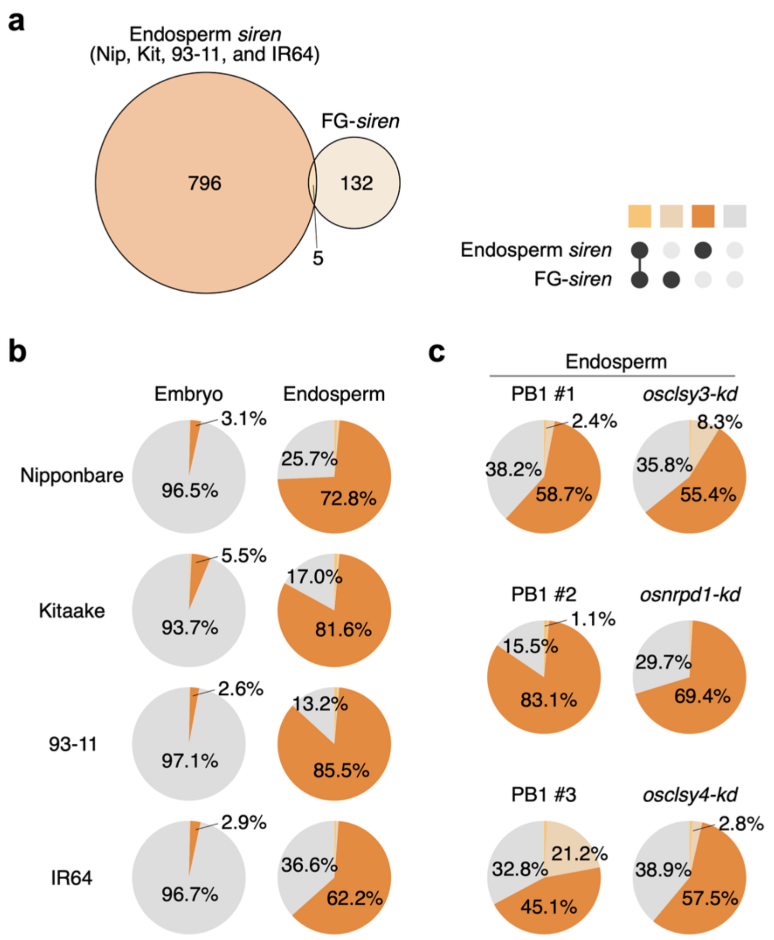


**Supplementary Fig. 7. Comparison of 24-nt siRNA accumulation at endosperm *siren* loci and FG-*siren* loci. a,** Overlap between endosperm *siren* loci and FG-*siren* loci. **b,c,** Relative abundance of 24-nt siRNAs expressed in each intersection category: embryo and endosperm from Nipponbare, Kitaake, 93-11, and IR64 varieties (**b**), and RdDM-related mutants and their corresponding wild types in the Pusa Basmati-1 (PB1) background (**c**). sRNA-seq datasets obtained from public repositories[1,2,10–12] and are listed in Supplementary Data 1. Genome coordinates for endosperm *siren* loci were obtained from a previous study[2], and comparisons with FG-*siren* loci are listed in Supplementary Data 8.

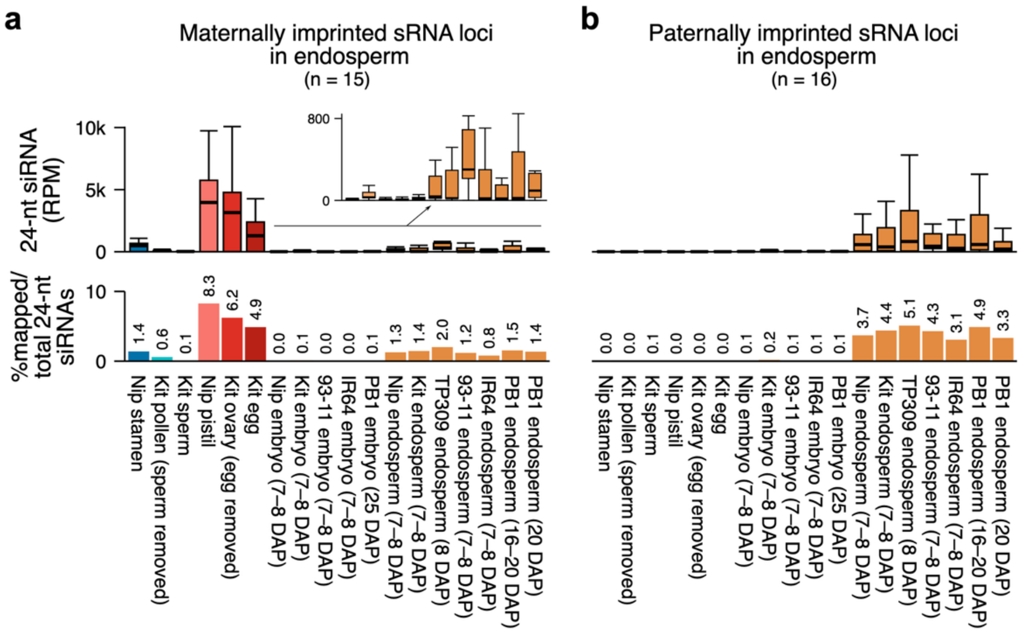


**Supplementary Fig. 8. Comparison of 24-nt siRNA accumulation at imprinted sRNA loci in endosperm. a,b,** 24-nt siRNA expression levels at maternally (**a**) and paternally (**b**) imprinted sRNA loci in endosperm across reproductive tissues from different rice varieties. In the lower panel, the relative abundances of 24-nt siRNAs derived from each sRNA loci are shown. sRNA-seq datasets obtained from public repositories[1–4,11,12,18,19] and are listed in Supplementary Data 1. Genome coordinates for endosperm imprinted sRNA loci were obtained from a previous study[1]. Abbreviations for rice cultivars: Nip, Nipponbare; Kit, Kitaake; TP309, Taipei 309; PB1, Pusa Basmati 1.

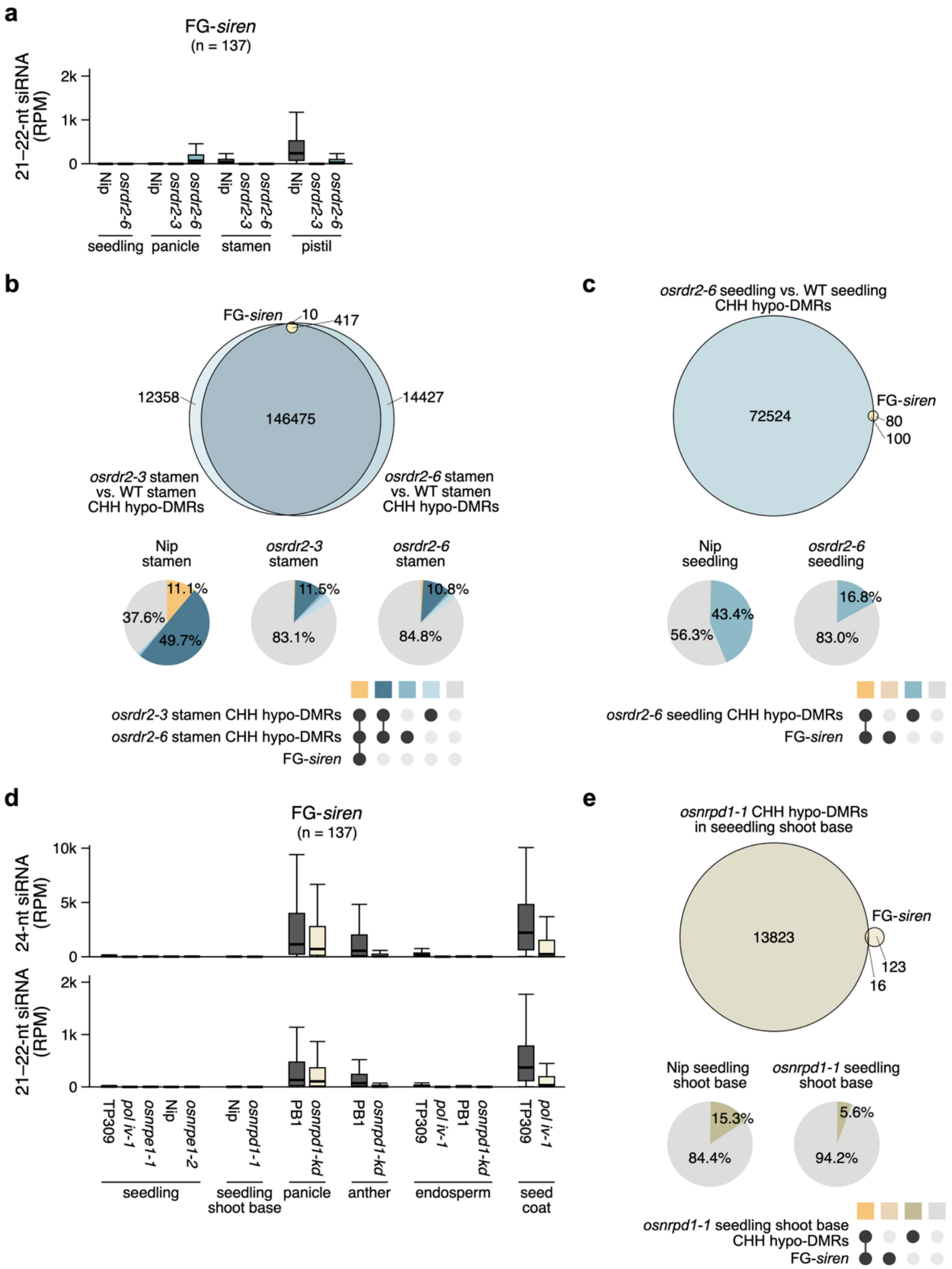


**Supplementary Fig. 9. Comparison of 24-nt and 21–22-nt siRNA accumulation at FG-*siren* loci in canonical RdDM-related mutants. a,** 21–22-nt siRNA expression at FG-*siren* loci in *osrdr2* mutants across vegetative and reproductive tissues. **b,c,** Comparison between FG-*siren* loci and *osrdr2* CHH hypo-DMRs in stamens (**b**) and seedlings (**c**) compared to wild type. **d,** 24-nt and 21–22-nt siRNA expression at FG-*siren* loci in Pol IV- and Pol V-related mutants across vegetative and reproductive tissues. **e,** Comparison between FG-*siren* loci and *osnrpd1-1* CHH hypo-DMRs in seedling shoot bases compared to wild type. For **b,c,e**, scaled Venn diagrams depict the relationships among the compared loci, and pie charts show the relative abundance of 24-nt siRNAs expressed in each intersection category. sRNA-seq datasets obtained from public repositories[10,17–20] and are listed in Supplementary Data 1. Genome coordinates for *osrdr2* CHH hypo-DMRs in stamens and seedlings compared to WT[18] and *osnrpd1* CHH hypo-DMRs in seedling shoot base compared to WT[17] were obtained from previous studies, and comparisons with FG-*siren* loci are listed in Supplementary Data 10.

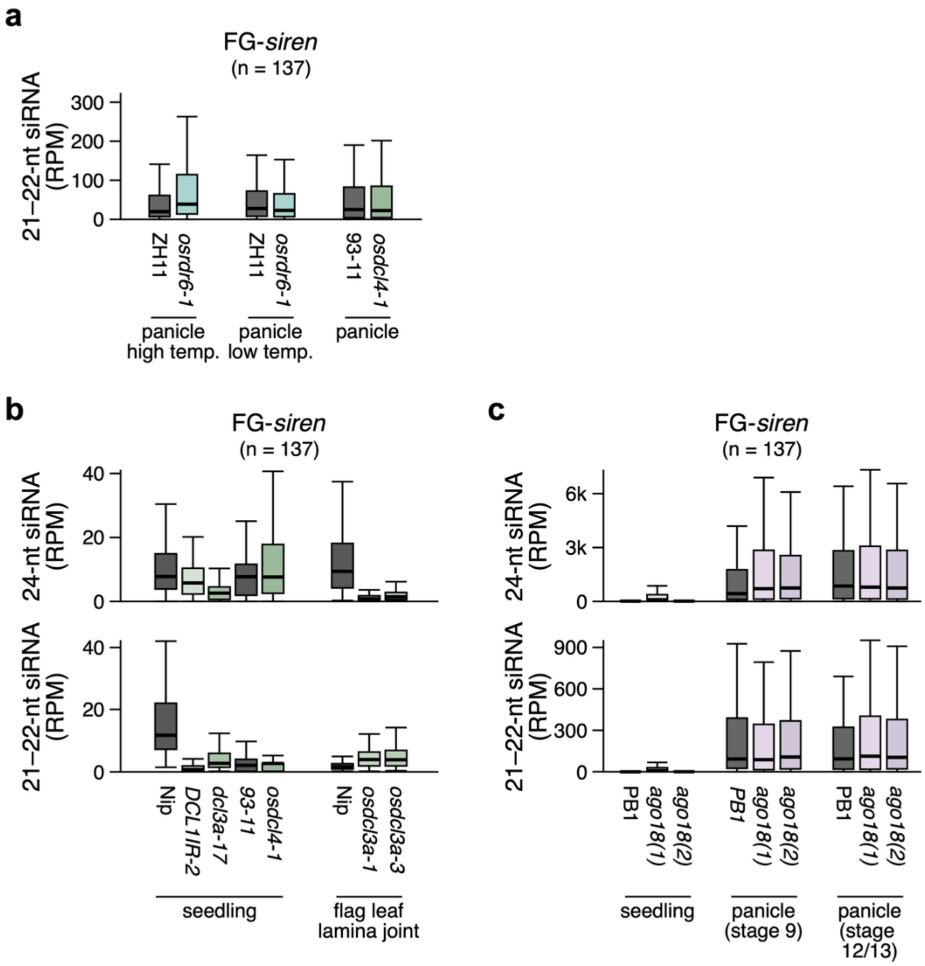


**Supplementary Fig. 10. Comparison of 24-nt and 21–22-nt siRNA accumulation at FG-*siren* loci in non-canonical RdDM-related mutants. a,** 21–22-nt siRNA expression at FG-*siren* loci in WT, *osrdr6-1*, and *osdcl4-1* in panicles. **b,c,** 24-nt and 21–22-nt siRNA expression at FG-*siren* loci in DCL-related (**b**) and AGO18-related (**c**) mutants across vegetative and reproductive tissues. sRNA-seq datasets obtained from public repositories[7–9,13,14] and are listed in Supplementary Data 1.

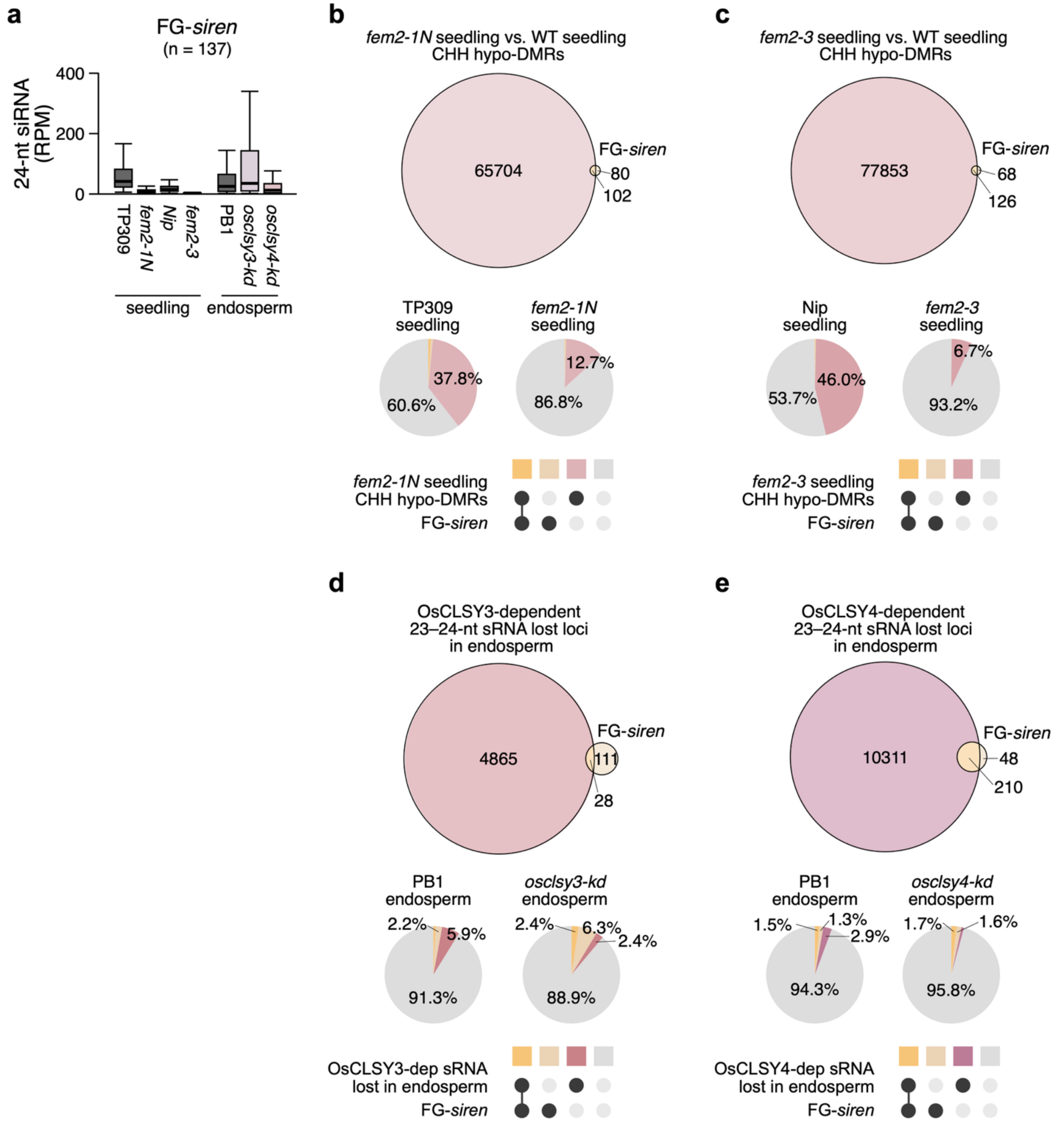


**Supplementary Fig. 11. Comparison of 24-nt siRNA accumulation at FG-*siren* loci in CLSY mutants. a,** 24-nt siRNA expression at FG-*siren* loci in CLSY mutants across vegetative and reproductive tissues. **b,c,** Comparison between FG-*siren* loci and CHH hypo-DMRs in *fem2-1N* (**b**) and *fem2-3* (**c**) seedlings compared to wild type. **d,e,** Comparison between FG-*siren* loci and OsCLSY3-dependent (**d**) and OsCLSY4-dependent (**e**) 23–24-nt sRNA loci lost in endosperm. For **b–e**, scaled Venn diagrams depict the relationships among the compared loci, and pie charts show the relative abundance of 24-nt siRNAs expressed in each intersection category. sRNA-seq datasets obtained from public repositories[11,12,18,21] and are listed in Supplementary Data 1. Genome coordinates for *fem2* CHH hypo-DMRs in seedlings compared to WT[18] and OsCLSY3/4-dependent 23–24-nt sRNA loci lost in endosperm[12] were obtained from previous studies, and comparisons with FG-*siren* loci are listed in Supplementary Data 9,10.

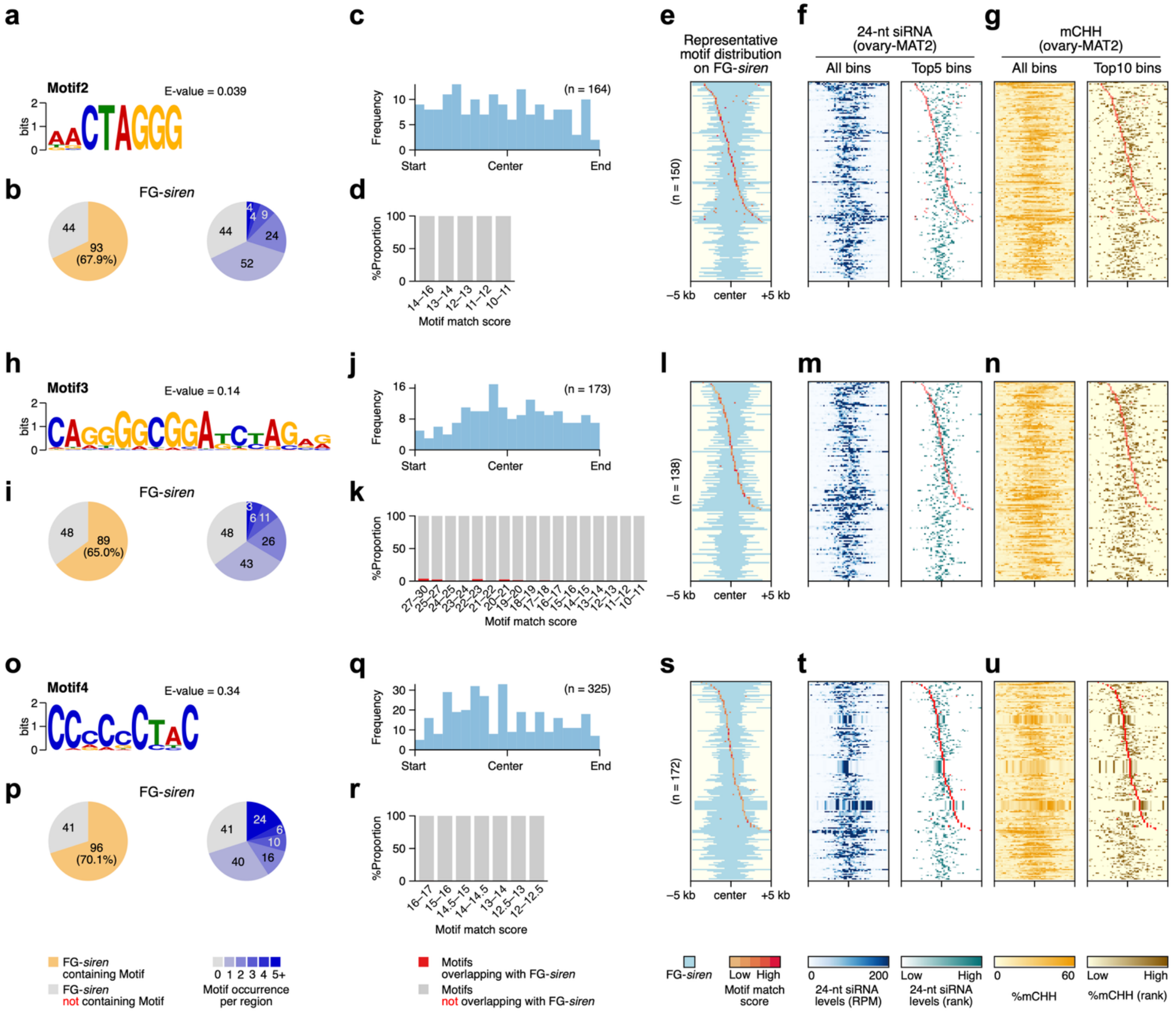


**Supplementary Fig. 12. DNA motifs enriched in FG-*siren* loci. a–u,** DNA motifs identified using STREME analysis: Motif2 (**a–g**), Motif3 (**h–n**), and Motif4 (**o–u**). **a,h,o,** DNA motif sequences enriched in FG-*siren* loci. **b,i,p,** Occurrence of DNA motifs at FG-*siren* loci. **c,j,q,** Distribution of DNA motifs across FG-*siren* loci in proportional bins (n = 20). **d,k,r,** Proportion of motifs overlapping with FG-*siren* loci across motif match score bins, where scores are defined by similarity to the consensus sequence. **e–g,l–n,s–u,** Motif distribution (**e,l,s**), 24-nt siRNA levels (**f,m,t**), and CHH methylation levels (**g,n,u**) across FG-*siren* loci. For each locus, motifs with the highest match score were selected as representative. The top five and top ten bins per locus were highlighted for 24-nt siRNA and CHH methylation levels, respectively. Information on Motif1 (named FRE) is shown in **Fig. 4**. More information on DNA motifs is available in Supplementary Data 11.

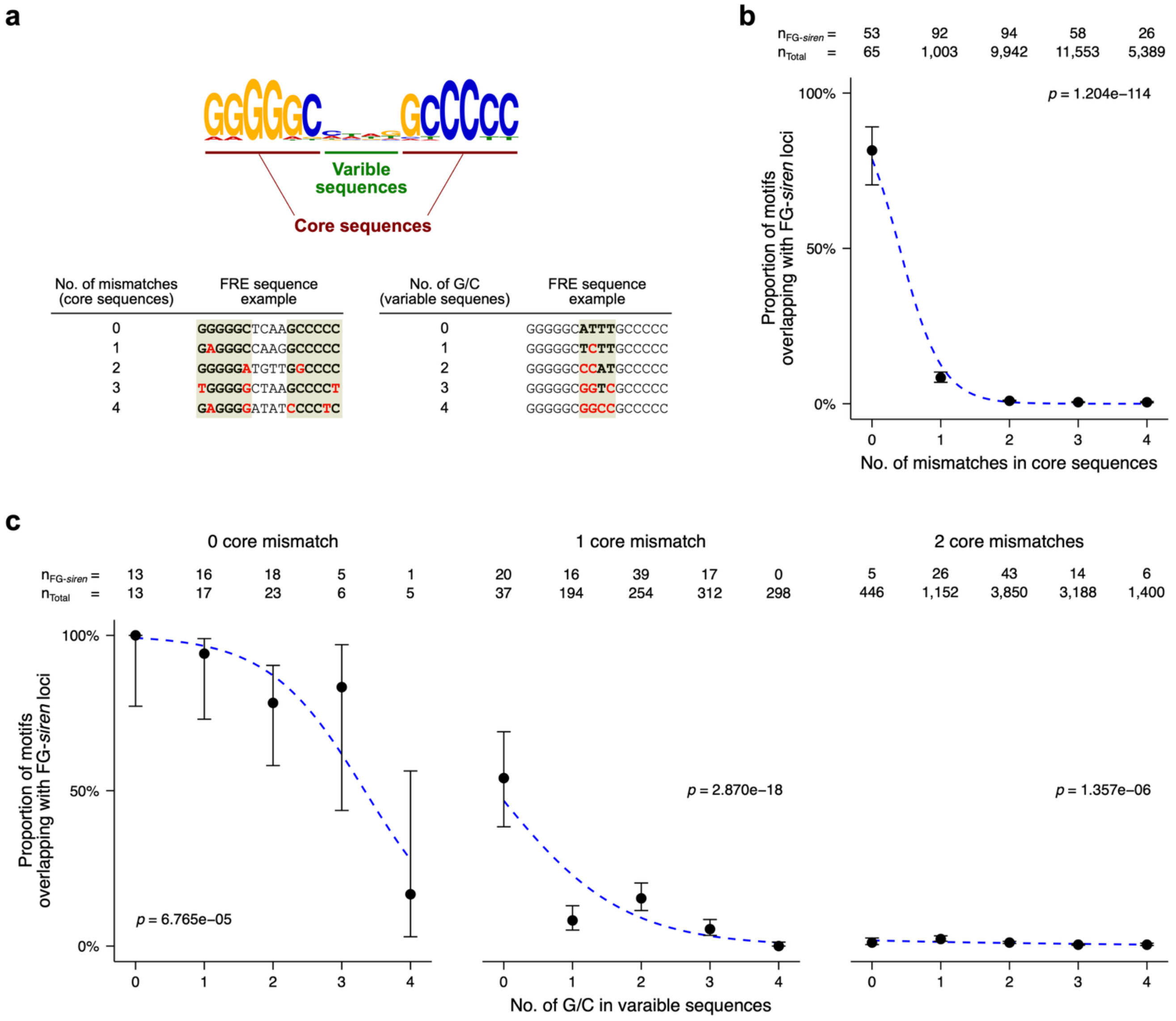


**Supplementary Fig. 13. Effects of base composition within FRE motifs on siRNA expression. a,** Core and variable sequences of the FRE motif. Examples of FRE sequences are shown according to the number of mismatches in the core sequences and the number of G/C bases in the variable sequences. **b,c,** Proportion of FRE motifs overlapping FG-*siren* loci according to the number of mismatches in the core sequences (**b**) and the number of G/C bases in the variable sequences (**c**). Error bars indicate 95% confidence intervals calculated using the Wilson method. Blue dashed lines were fitted using a generalized linear model with a quasibinomial error distribution. The counts of total FRE motifs and FG-*siren*-overlapping FRE motifs are indicated. Statistical significance for monotonic trends in the proportion of FRE motifs overlapping FG-*siren* loci across ordered mismatch or G/C content categories was determined using the Cochran-Armitage trend test.

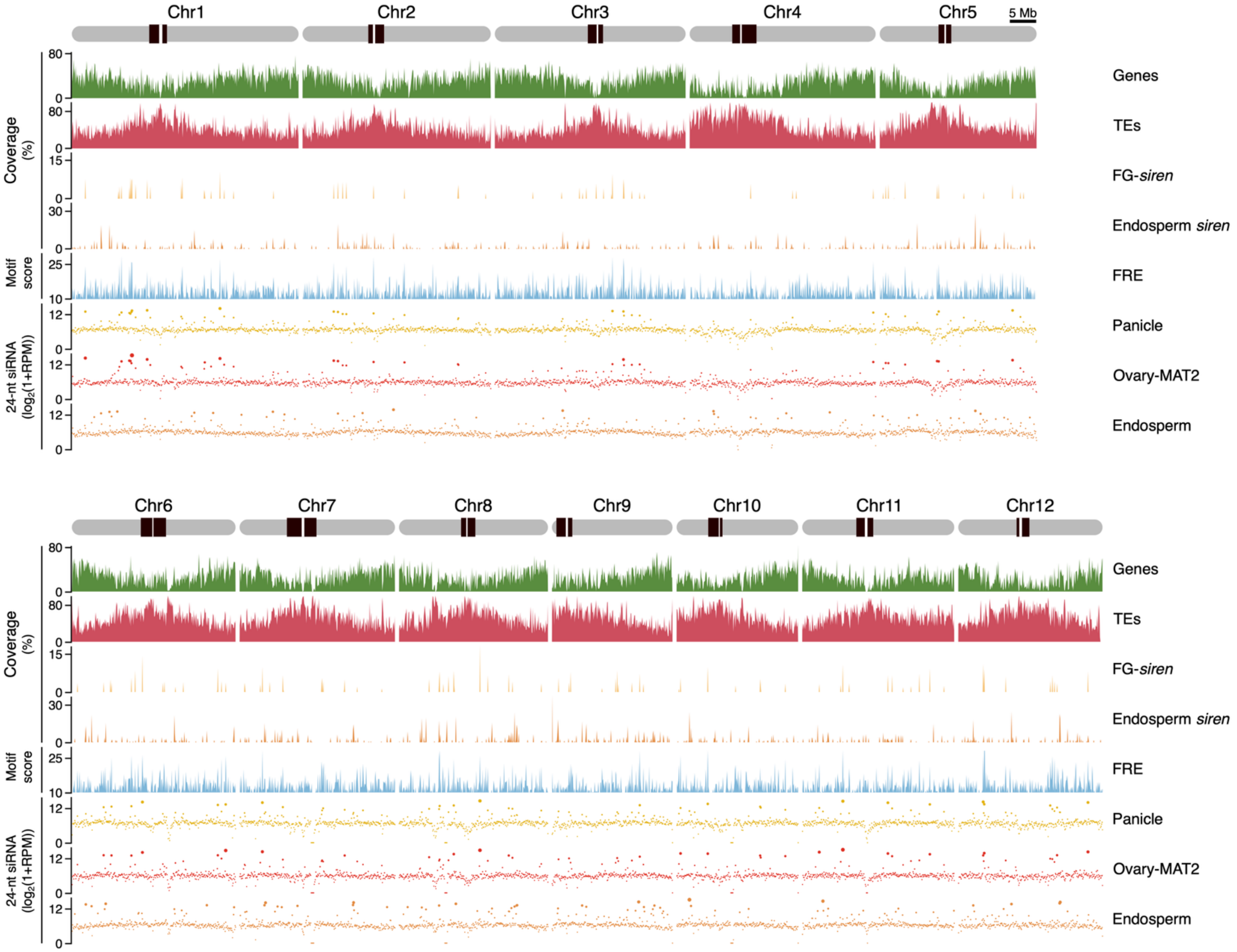


**Supplementary Fig. 14. Genome-wide distribution of FG-*siren* loci and FRE motifs.** Chromosomal distribution of genes, TEs, FG-*siren* loci, endosperm *siren* loci, FRE motifs, and 24-nt siRNA levels in Nipponbare panicle, ovary (MAT2 stage), and endosperm along all rice chromosomes, summarized in 100-kb bins. Pericentromeric heterochromatin was defined based on the genetic recombination-suppressed regions of Tian et al. (2009)[24]; the remaining regions are considered chromosomal arms. sRNA-seq datasets for Nipponbare panicle and endosperm were obtained from public repositories[1,18] and are listed in Supplementary Data 1. Genome coordinates for endosperm *siren* loci were obtained from a previous study[2].

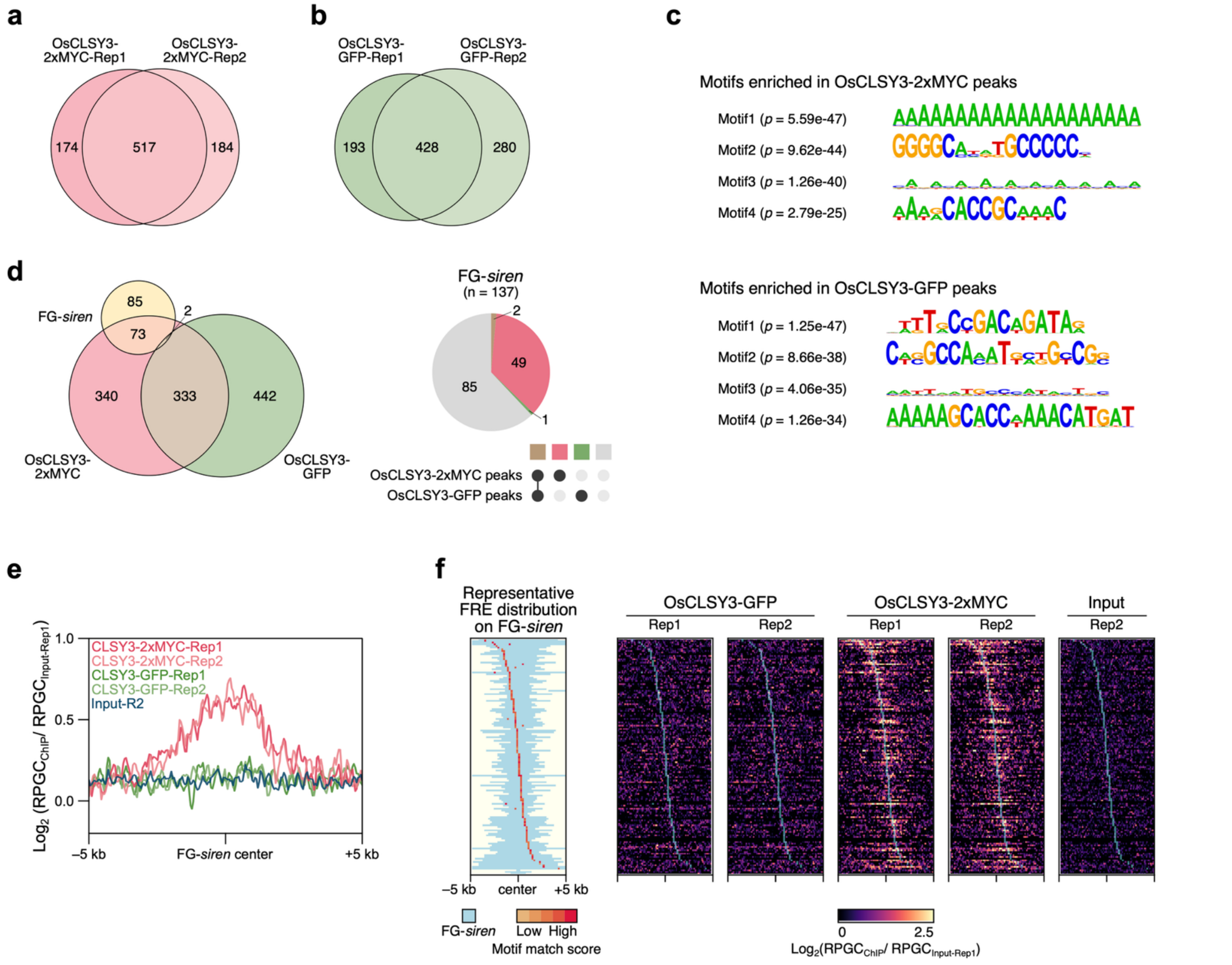


**Supplementary Fig. 15. ChIP-seq analysis of OsCLSY3. a,b,** Relationships between replicates of ChIP data from OsCLSY3-2xMYC (**a**) and OsCLSY3-GFP (**b**). Identified ChIP peaks are listed in Supplementary Data 14. **c,** DNA motifs enriched in ChIP peaks from OsCLSY3-2xMYC and OsCLSY3-GFP. Identified motifs are listed in Supplementary Data 15. **d,** Relationships among FG-*siren* loci and ChIP peaks from OsCLSY3-2xMYC and OsCLSY3-GFP (left). Proportion of FG-*siren* loci overlapping with ChIP peaks (right). Detailed information is available in Supplementary Data 16. **e,** Profile plots showing average ChIP enrichment across FG-*siren* loci. **f,** Heatmaps showing OsCLSY3 ChIP enrichment across FG-*siren* loci. For each locus, motifs with the highest match score were selected as representative.

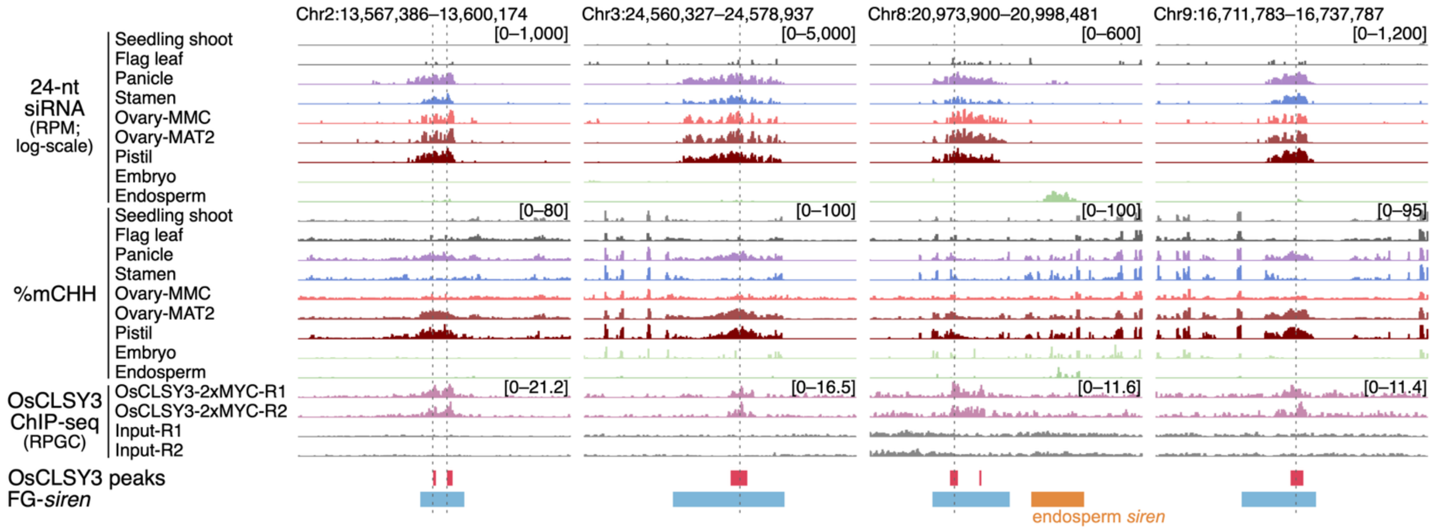


**Supplementary Fig. 16. siRNA landscape, DNA methylation, and OsCLSY3 enrichment at representative FG-*siren* loci.** 24-nt siRNA and CHH methylation levels across rice tissues, together with OsCLSY3 enrichment in rice panicles, at four representative FG-*siren* loci. Adjacent endosperm *siren* loci are shown in orange. Grey dotted lines represent the FRE motifs. Coordinates for endosperm *siren* loci were obtained from Rodrigues et al. (2021)[2]. OsCLSY3 ChIP-seq datasets were obtained from public repositories[11] and are listed in Supplementary Data 13.

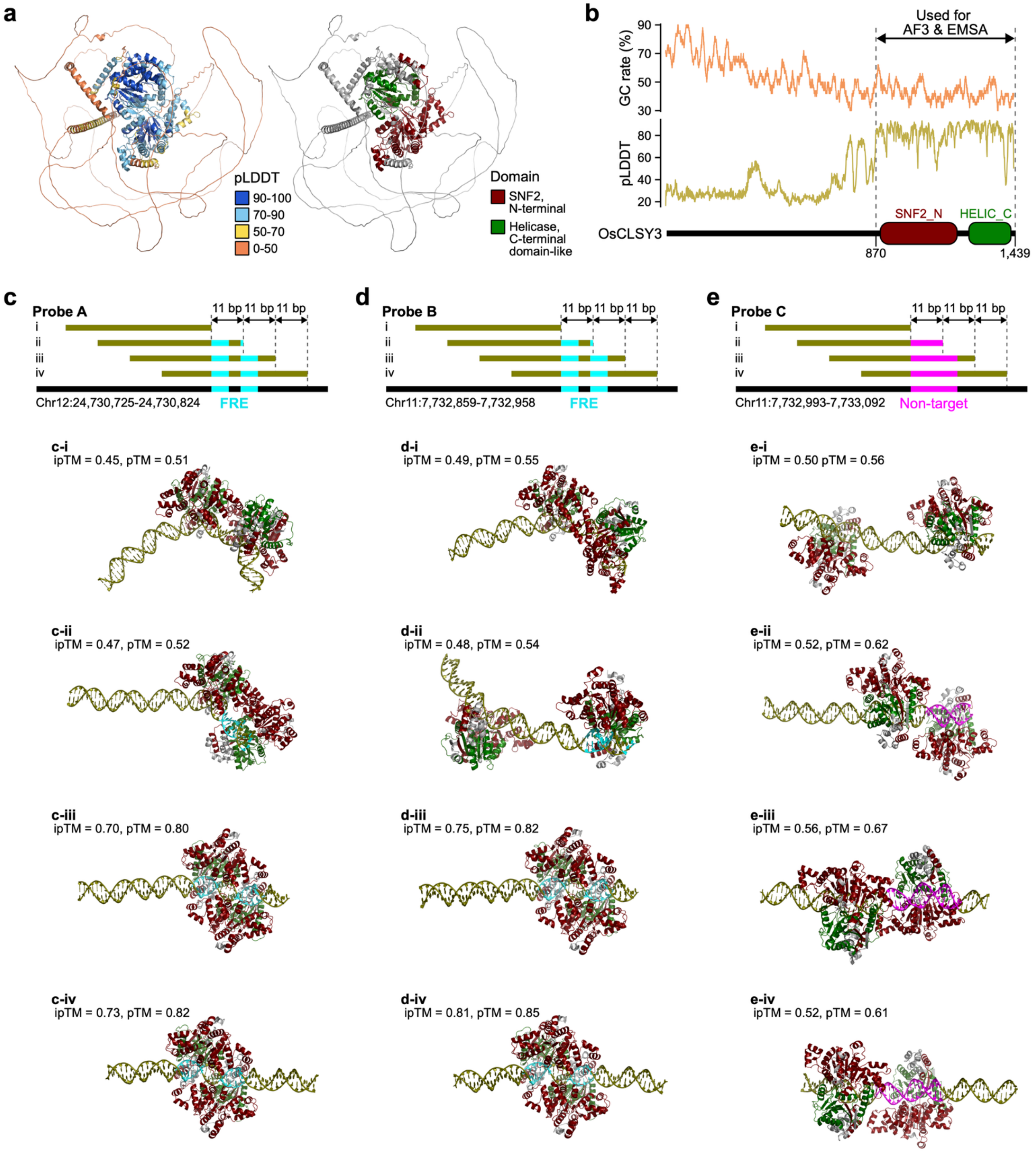


**Supplementary Fig. 17. Predicted interaction between OsCLSY3 and the FRE motif using AlphaFold3. a,** AlphaFold3 (AF3)-predicted structure of OsCLSY3, colored by pLDDT (prediction confidence; left panel) and by functional domains (right panel). **b,** Schematic representation of OsCLSY3 showing GC content (25-bp sliding window) and per-residue pLDDT scores from AF3. **c–e,** AF3 predictions for OsCLSY3 bound to two FRE-containing probes (Probe A (**c**) and Probe B (**d**)) and a non-FRE control (Probe C (**e**)). pTM (predicted template modeling) and ipTM (interface pTM) scores indicate the confidence in the predicted complex fold and the relative positioning of subunits within the complex, respectively. Probes used for AF3 prediction are listed in Supplementary Data 17.

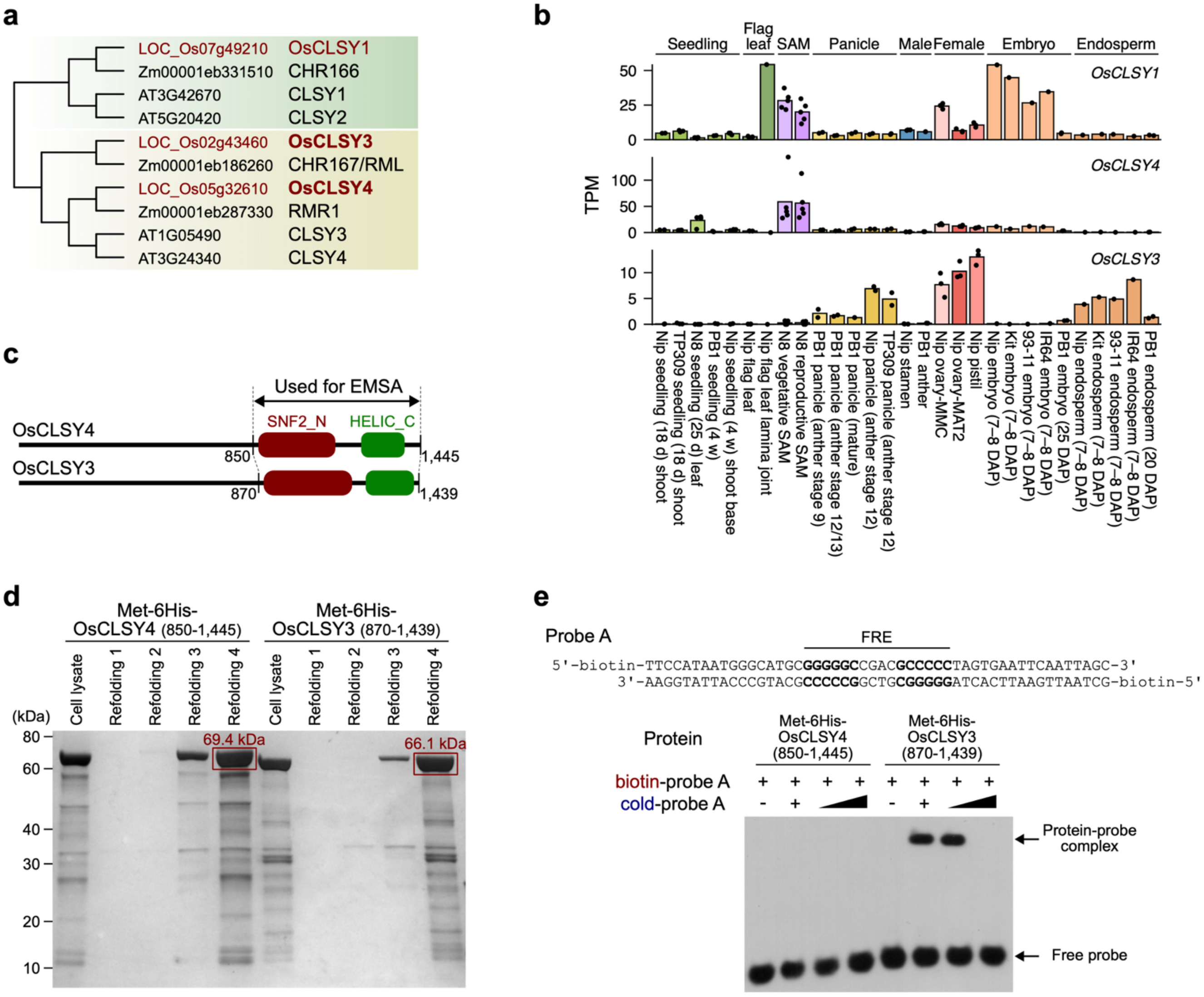


| Buffer | Components |
|---|---|
| Refolding1 | 50 mM Tris-HCl, 150 mM NaCl, 10% Glycerol, pH 8.0 |
| Refolding2 | 50 mM Tris-HCl, 500 mM NaCl, 10% Glycerol, pH 8.0 |
| Refolding3 | 50 mM Tris-HCl, 150 mM NaCl, 0.5 M L-Arginine, 10% Glycerol, pH 8.0 |
| Refolding4 | 50 mM Tris-HCl, 150 mM NaCl, 1.0 M L-Arginine, 10% Glycerol, pH 8.0 |

**Supplementary Fig. 18. Validation of direct interaction between OsCLSY3 and the FRE motif. a,** Phylogenetic analysis of CLSY genes in *Arabidopsis,* maize, and rice. Schematic topology is based on Trujillo et al. (2018)[25] and Pal et al. (2024)[11]. **b,** Transcript levels of OsCLSY genes across vegetative and reproductive tissues. mRNA-seq datasets for Nipponbare panicle and endosperm were obtained from public repositories[2,5,9–11,14,17–19] and are listed in Supplementary Data 3. **c,** Schematic representation of OsCLSY proteins used for EMSA. **d,** Purification of 6His-tagged truncated OsCLSY4 and OsCLSY3 proteins. Proteins purified using refolding buffer 4 were used for EMSA. **e,** EMSA showing interactions of OsCLSY3, but not OsCLSY4, with the FRE-containing probe. Cold probe gradients represent 3- and 30-fold excess over biotin-labeled probes. Probe sequences were determined based on the genomic sequences of the representative FG-siren loci in Fig. 5f. Probes used for EMSA are listed in Supplementary Data 17.

### Supplementary Notes

**Supplementary Note 1.** Analysis of OsCLSY3 ChIP-seq.

The enrichment of FRE enriched in FG-*siren* loci (**Fig. 4**) and the strong positive correlation between OsCLSY3 transcript levels and 24-nt siRNA levels at FG-siren loci (**Fig. 5a**) led us to test whether OsCLSY3 associates with FRE in vivo. Pal et al. (2024) performed ChIP-seq in panicle tissues from transgenic rice expressing OsCLSY3 driven by its native promoter and tagged at the C terminus (GFP and 2xMYC)[11]. We re-analyzed these ChIP-seq datasets.

*Rationale for re-analysis*

We first inspected the bigWig signal tracks deposited with the original study (hereafter "as-provided"; GSE229961). The patterns shown in the original figures could be reproduced when visualizing input-subtracted difference tracks generated by direct subtraction using a single input control (Input-R1) (**Supplementary Note Fig. 1a,c,e**). However, inspection of the as-provided tracks without subtraction showed that apparent ChIP enrichments at the reported OsCLSY3 peak regions frequently coincide with strong input signal (**Supplementary Note Fig. 1b,d,f**), suggesting that the displayed patterns are highly dependent on the subtraction step and input track structure. We therefore re-analyzed the dataset starting from the raw reads, following the workflow described by Pal et al. (2024)[11].

*Re-mapping OsCLSY3 ChIP-seq reads*

The original publication[11] reported that 100-bp paired-end ChIP-seq reads were aligned to the IRGSP1.0 genome[26] using Bowtie1 with -v 3 -k 1 -y -a --best --strata[27]. Because -a reports all valid alignments whereas -k 1 reports up to one valid alignment, these options imply different reporting behaviors. We therefore tested two implementations: Bowtie1 with -a (*bowtie1_a*) and Bowtie1 with -k 1 (*bowtie1_k1*), while keeping the remaining parameters unchanged. In parallel, we also aligned reads with Bowtie2 using --very-sensitive-local -N 1 (*bowtie2_N1_local*), as Bowtie2 is generally recommend for reads longer than 50 bp[28] (**Supplementary Data 13**).

Comparison among these three mapping strategies showed that the ChIP/Input enrichment patterns observed in the as-provided signal tracks (**Supplementary Note Fig. 1**) were reproduced using *bowtie1_a*. In contrast, signal at most originally reported OsCLSY3 peak regions was reduced or absent when using *bowtie1_k1* or *bowtie2_N1_local* (**Supplementary Note Fig. 2a,b**). To further assess whether multi-mapping reads contributed the original identification of OsCLSY3 ChIP peaks, we examined the number of reported alignments for reads mapped to these peaks. Under *bowtie1_a*, reads mapped to the reported ChIP peaks had an average of ~16.6 alignments, substantially higher than the chromosome-wide average (~4.4). Moreover, reads containing DNA motifs enriched in OsCLSY3 ChIP peaks showed even higher numbers of alignments (~46.6

for motif 1 to ~29.2 for motif 4) (**Supplementary Note Fig. 2a,c,d**). Together, these results suggest that the reported OsCLSY3 ChIP peaks and associated motif enrichments by Pal et al. (2024)[11] are highly sensitive to multi-mapping reads and may therefore include regions that do not robustly reflect OsCLSY3-specific enrichment. Accordingly, we used *bowtie2_N1_local* for downstream analyses to obtain a more conservative alignment approach for peak calling.

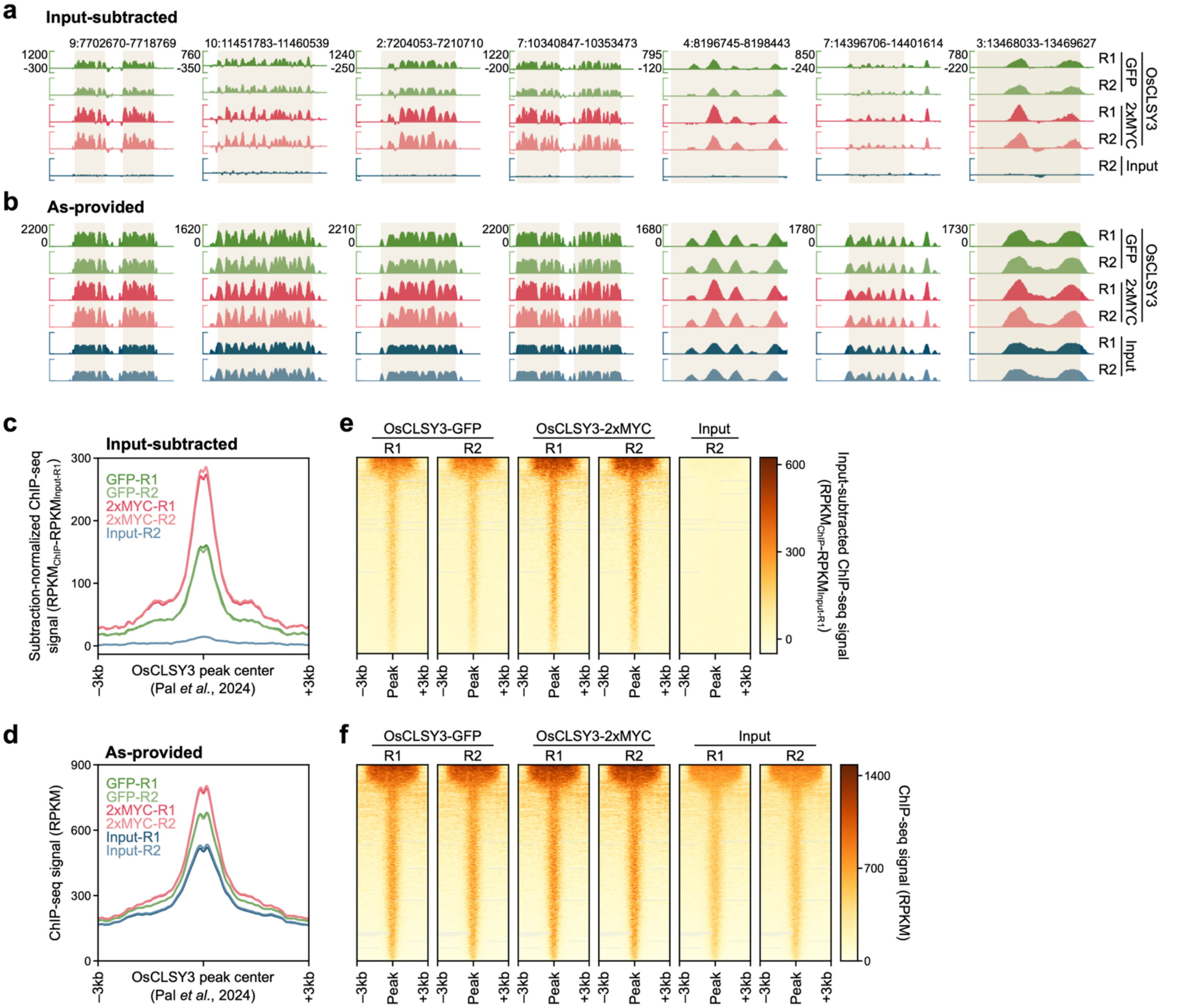


**Supplementary Note Fig. 1. Inspection of signal track representations for OsCLSY3 ChIP-seq. a,b,** Genome browser screenshots displaying Input-subtracted tracks (**a**) and as-provided ChIP tracks (**b**) at representative OsCLSY3 ChIP peak regions. **c–f,** Profile plots (**c,d**) and heatmaps (**e,f**) showing Input-subtracted track (**c,e**) and as-provided ChIP signals (**d,f**) within ±3 kb flanking regions from OsCLSY3 ChIP peak centers. Peak coordinates are according to the original publication[11].

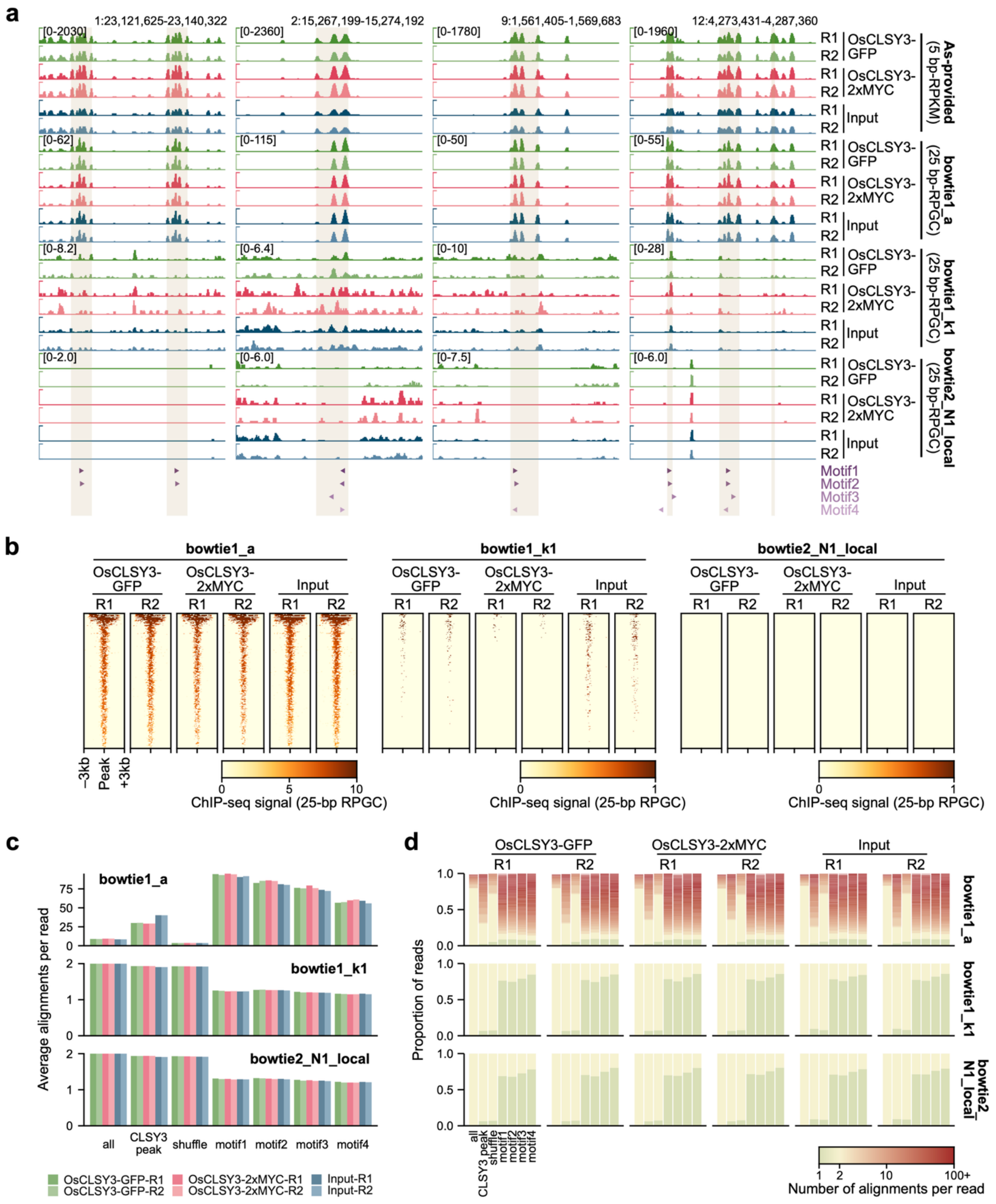


**Supplementary Note Fig. 2. Re-mapping OsCLSY3 ChIP-seq reads. a,** Genome browser screenshots displaying ChIP signal tracks at representative OsCLSY3 ChIP peak regions. Shaded regions and arrowheads indicate OsCLSY3 ChIP peaks and enriched DNA motifs (motif 1–4). Peak coordinates and motif annotations are according to the original publication[11]. **b,** Heatmaps showing ChIP signals within ±3 kb flanking regions from OsCLSY3 ChIP peak centers. **c,d,** Average number of alignments per read (**c**) and the proportion of reads stratified by the number of alignments per read (**d**) for reads mapped to different genomic categories (all reads, OsCLSY3 peaks, and shuffled regions derived from OsCLSY3 peaks) and for reads containing motif 1–4. For **a–d**, results from *bowtie1_a*, *bowtie1_k1*, and *bowtie2_N1_local* are shown.

**Supplementary Data** (available as **Additional Files**)

**Supplementary Data 1.** Basic statistics of sRNA-seq datasets.

**Supplementary Data 2.** Basic statistics of EM-seq datasets.

**Supplementary Data 3.** Basic statistics of mRNA-seq datasets.

**Supplementary Data 4.** List of 24-nt siRNA abundant loci in developing ovaries and flag leaves.

**Supplementary Data 5.** List of FG-*siren* loci.

**Supplementary Data 6.** Comparison of FG-*siren* loci with 24-nt siRNA abundant loci in flag leaf.

**Supplementary Data 7.** Comparison of FG-*siren* loci with 24-nt siRNA clusters across vegetative and reproductive tissues.

**Supplementary Data 8.** Comparison of FG-*siren* loci with CHH hyper-DMRs in reproductive tissues compared to seedlings.

**Supplementary Data 9.** Comparison of FG-*siren* loci with endosperm sRNA loci.

**Supplementary Data 10.** Comparison of FG-*siren* loci with CHH hypo-DMRs in RdDM-related mutants.

**Supplementary Data 11.** List of DNA motifs enriched in FG-*siren* loci.

**Supplementary Data 12.** Pearson correlation between 24-nt siRNA levels at FG-*siren* loci and transcript levels of all expressed genes in rice tissues.

**Supplementary Data 13.** Basic statistics of ChIP-seq datasets.

**Supplementary Data 14.** List of peaks obtained from OsCLSY3 ChIP-seq analysis.

**Supplementary Data 15.** List of DNA motifs enriched in OsCLSY3 ChIP peaks.

**Supplementary Data 16.** Comparison of FG-*siren* loci with OsCLSY3 ChIP peaks.

**Supplementary Data 17.** Oligonucleotides used in this study.
